# Transfer learning across molecular graphs for predicting protein-ligand affinities and their changes upon mutations

**DOI:** 10.1101/2025.07.19.665665

**Authors:** Yunzhuo Zhou, YooChan Myung, Alex G. C. de Sá, David B. Ascher

## Abstract

Predicting protein-ligand binding affinity and mutation-induced affinity changes (ΔΔ*G*) remains challenging due to limited data and complex interaction mechanisms. Here we present DDMuffin, a deep learning framework that integrates structural, sequence, and interaction graph features, employing transfer learning and stringent dataset partitioning to achieve reliable generalization. DDMuffin demonstrates strong predictive accuracy on the rigorous LP-PDBBind benchmark (Pearson *r* up to 0.70 after excluding top 10% outliers; RMSE = 1.48 kcal mol⁻¹). In evaluating mutation-induced affinity changes for clinically relevant kinase inhibitors, DDMuffin achieves competitive average performance (mean Spearman *ρ* = 0.39), outperforming or matching several established methods. The approach provides valuable insights into mutation-driven drug resistance and interpretable ligand binding mechanisms, particularly advantageous for guiding inhibitor design and personalized therapeutic strategies. To facilitate broad application, we deployed DDMuffin as an accessible web server at https://biosig.lab.uq.edu.au/ddmuffin/, promoting systematic exploration of protein-ligand interactions in drug discovery research.

## Introduction

Protein-ligand interactions are fundamental to a wide range of biological processes and form a cornerstone of structure-based drug design. Accurate prediction of binding affinity is crucial for drug screening and lead optimization in pharmacology^1^. However, missense mutations in proteins can dramatically alter ligand binding affinity, leading to genetic diseases and undermining the efficacy of therapeutic drugs via drug resistance^2^. In cancer and infectious diseases, for example, acquired mutations in drug targets are a major mechanism of resistance that often causes treatments to fail^3,4^. These challenges highlight the importance of understanding and predicting how protein-ligand interactions respond to genetic variations.

Advances in DNA sequencing have uncovered an enormous number of protein variants across clinical isolates and patient samples^5^, yet determining which mutations will affect ligand binding or drug efficacy remains difficult^6^. Experimental techniques can measure mutation-induced changes in binding free energy (ΔΔ*G*) with high accuracy^7^, but such assays are time-consuming and costly, making it infeasible to survey the vast mutational space comprehensively. This creates an urgent need for reliable computational methods to predict the effects of mutations on protein-ligand binding. Effective *in silico* prediction of affinity changes would help identify pathogenic or drug-resistant mutations and guide the development of new therapeutic strategies.

Computational methods, particularly deep learning models have recently achieved impressive results in predicting protein-ligand binding affinities from large datasets^1,8–15^. Meanwhile, specialized computational methods can estimate the effects of point mutations on protein properties, including stability^16–18^ and interactions^19–28^. However, these two problem domains, binding affinity prediction and mutation impact prediction, have largely been addressed in isolation. Integrating them is essential to build more powerful and generalizable predictors that can handle both wild-type and mutant protein-ligand complexes.

Developing robust predictive models requires strict evaluation. However, current benchmarking practices often suffer from data leakage, where structurally or chemically similar complexes unintentionally appear in both training and test datasets, inflating apparent performance^29^. For instance, widely-used benchmarks like PDBbind^30^ contain significant overlap between training and test splits, leading to overly optimistic assessments. To address this issue, rigorously non-redundant datasets such as leak-proof (LP)-PDBBind^29^ have been developed, ensuring unbiased evaluations and more realistic performance estimates.

Here we present DDMuffin, a deep learning framework for predicting protein-ligand binding affinities and their changes upon mutation, emphasizing robust generalization through careful design and validation. In designing DDMuffin, we employed strict similarity-based dataset partitioning similar to LP-PDBBind^29^, ensuring that no protein in the test set has close sequence homology to any training protein, and that no ligand scaffold is shared between training and test complexes. Furthermore, we validated our model using completely independent external datasets to assess its predictive power on novel targets. This strategy provides confidence that DDMuffin’s performance reflects true predictive ability rather than exploitation of overlap or biases in the data.

DDMuffin leverages transfer learning by first pre-training on an extensive dataset of protein-ligand complexes, capturing diverse structural and chemical interaction features, and is subsequently fine-tuned using mutation-specific data to precisely estimate ΔΔ*G*. Its architecture leverages graph convolutional networks (GCNs) to integrate multi-scale molecular representations, including protein graph with residue-level ProtT5 embeddings^31^, ligand graph with pharmacophoric features, and protein-ligand interaction graph with atomic-level interaction fingerprints derived from Arpeggio^32^. Crucially, DDMuffin bridges absolute and relative affinity prediction within one framework, providing researchers with a one-stop tool for evaluating both a compound’s baseline binding strength and how that binding might shift due to target mutations. To maximize accessibility, we have deployed DDMuffin as a user-friendly web server at https://biosig.lab.uq.edu.au/ddmuffin/, allowing experimental and computational scientists to easily assess binding affinities and ΔΔ*G* for their proteins of interest without the need for extensive computational resources. This platform is expected to aid both fundamental and translational research, for example, by helping to anticipate resistance mutations, prioritize variant testing, or design more robust inhibitors, ultimately facilitating progress in structural biology and drug discovery.

## Results

### Overview of the DDMuffin Method

DDMuffin is a computational approach designed to predict how missense mutations affect protein-ligand affinity (ΔΔ*G*). It adopts a two-stage deep learning strategy consisting of an initial pre-training step followed by a specialized fine-tuning phase. During pre-training, DDMuffin captures broadly applicable interaction patterns using the extensive Leak Proof PDBBind (LP-PDBBind) dataset^29^, which encompasses diverse protein-ligand complexes with experimentally measured affinities (IC_50_, *K*_d_ and *K*_i_) (**Fig. 1a**, left; **Supplementary Table 1**). The model’s capacity to generalize beyond its training dataset was validated independently using datasets involving SARS-CoV-2 main protease (Mpro)^33^, epidermal growth factor receptor (EGFR)^34^ complexes, and the BDB2020+^29^ dataset (compiled from recent entries in BindingDB^35^ deposited after 2020), demonstrating effective transferability even to structurally distinct proteins.

**Fig. 1.**
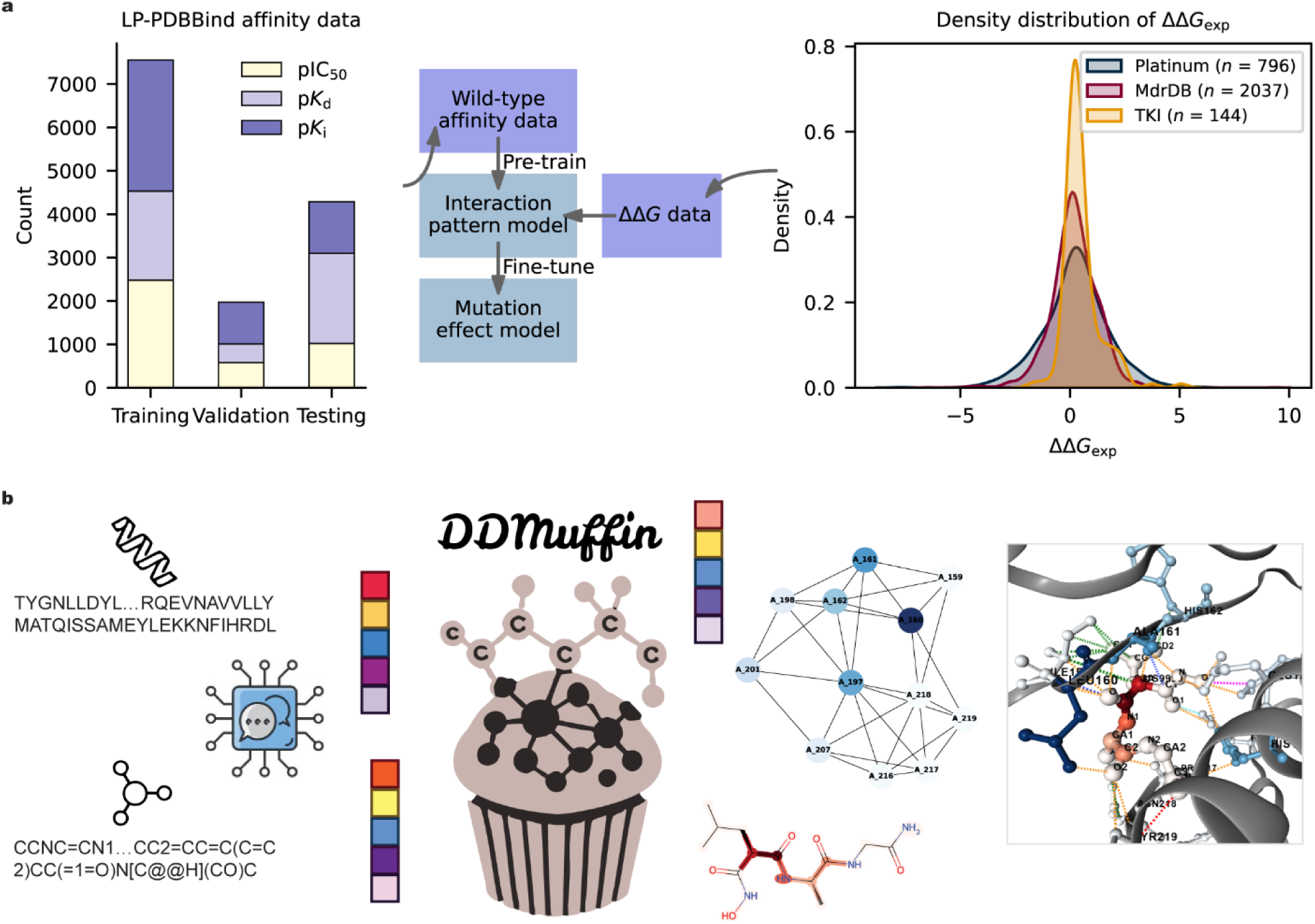
Overview of the DDMuffin (ΔΔ*G*: Mutation effects on protein-ligand affinity) Method. **a)** Data distribution and workflow. The bar chart (left) depicts the distribution of affinity measurement types (IC_50_, *K*_d_, *K*_i_) across the training, validation, and testing subsets within the LP-PDBBind dataset^29^ (**Supplementary Table 1**). The workflow diagram (centre) summarizes the two-step DDMuffin training approach: initial pre-training to model general protein-ligand interaction patterns, followed by targeted fine-tuning to specifically predict mutation-induced ΔΔ*G*. The density plot (right) illustrates the distribution of experimentally measured ΔΔ*G* values across independent datasets, including Platinum^36^, tyrosine kinase inhibitors (TKIs) interacting with ABL1 kinase^24^, and mutation-induced drug resistance database (MdrDB)^37^. **b)** Model architecture. DDMuffin integrates structural and biochemical insights through a modular deep learning framework combining linear feature-extraction layers with Graph Convolutional Networks. Molecular information is encoded via three complementary graphs: residue-level protein graphs annotated using ProtT5 embeddings^31^ to capture residue context, ligand atom-level graphs enriched with atomic pharmacophoric features, and protein-ligand interaction graphs annotated with pharmacophore types and Arpeggio-derived^32^ interatomic interaction features (**Methods**).

The fine-tuning phase leverages the specialized Platinum^36^ dataset, explicitly developed to capture mutation-induced affinity alterations. The fine-tuned model’s performance was further validated using independent datasets, including tyrosine kinase inhibitors (TKIs) binding to ABL1 kinase^24^ and the mutation-induced drug resistance database (MdrDB)^37^ (**Fig. 1a**, right). These validations demonstrated DDMuffin’s robust ability to predict mutation-induced affinity changes accurately.

DDMuffin integrates biochemical and structural information through linear feature-extraction layers coupled with GCNs (**Methods, Supplementary Fig. 1**). Molecular structures are represented via three complementary graph modalities: residue-level protein graphs annotated with ProtT5^31^ embeddings, ligand atom-level graphs enriched by atomic pharmacophoric features, and protein-ligand interaction graphs incorporating both pharmacophoric features and interatomic interactions derived from Arpeggio^32^ (**Fig. 1b**). ProtT5^31^ embeddings were specifically selected for their proven effectiveness in encoding contextual residue-level information, while Arpeggio-derived^32^ interactions were incorporated due to their precise characterization of critical interatomic contacts essential for accurate affinity predictions. By merging these multi-scale molecular representations, DDMuffin robustly predicted affinity changes caused by mutations, positioning it as a valuable computational tool for drug discovery and personalized therapeutic interventions.

### DDMuffin pre-training elucidates structural elements and feature contributions essential for predicting protein-ligand affinities

We pre-trained DDMuffin on the LP-PDBBind^29^ dataset to systematically uncover generalizable protein-ligand interaction patterns. To maintain data integrity and minimize redundancy, we preserved the original non-redundant LP-PDBBind training-validation-testing split. Additionally, the training set was partitioned into ten distinct folds based on protein^38^ and ligand^39^ similarities to facilitate rigorous hyperparameter optimization and ensure robust model generalization (**Methods**).

DDMuffin demonstrated strong predictive performance during pre-training, achieving Pearson correlation coefficients (*r*) of 0.58 (cross-validation on training), 0.60 (validation), and 0.54 (test) across the LP-PDBBind dataset (**Fig. 2a**, black metrics). Excluding the top 10% of predictions with the highest deviations from the regression fit significantly improved these correlations to 0.70, 0.68, and 0.66, respectively (**Fig. 2a**, red metrics). This suggests that DDMuffin reliably captures general binding-affinity trends but faces difficulties with specific challenging cases, notably involving lyase complexes, which were disproportionately represented among prediction outliers (**Supplementary Table 2**).

**Fig. 2.**
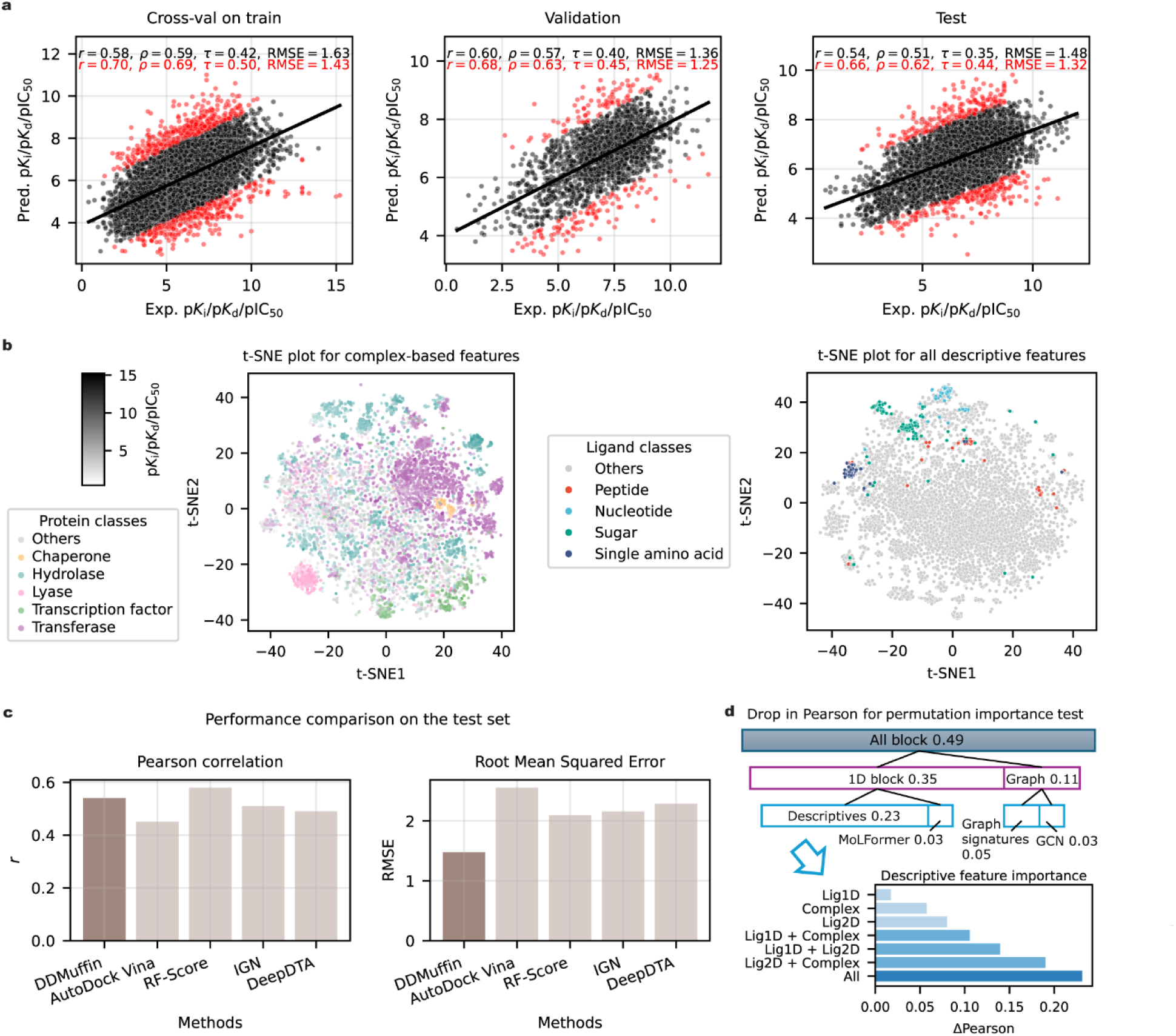
Pre-training affinity performance and feature importance analysis of DDMuffin using LP-PDBBind. **a)** Scatter plots comparing predicted versus experimental affinities (p*K*_i_, p*K*_d_, pIC_50_) for the LP-PDBBind^29^ dataset across training, validation and test sets. Pearson (*r*), Spearman (*ρ*), Kendall (*τ*) correlations, and Root Mean Squared Error (RMSE) values are shown at the top; black indicates results on the full dataset, red indicates performance after excluding the top 10% prediction outliers (red points), defined by greatest deviation from regression fits. **b)** t-SNE visualizations of descriptive features. Left panel: structural features derived solely from protein-ligand complexes, coloured by protein class and shaded by affinity strength. Right panel: all descriptive features, coloured by ligand class. **c)** Benchmark comparison of DDMuffin versus AutoDock Vina^8^, RF-Score^9^, InteractionGraphNet (IGN)^10^, and DeepDTA^11^ on the LP-PDBBind test set, comparing Pearson correlation (left) and RMSE (right) (**Supplementary Table 4**). Performance of benchmarking methods were directly taken from the LP-PDBBind paper^29^. **d)** Permutation importance analysis highlighting contributions of individual architecture blocks (top) and descriptive features (bottom) to model performance, measured by reductions in Pearson correlation upon permutation (**Methods**).

Performance assessment by affinity measurement type (pIC_50_, p*K*_d_ and p*K*_i_) revealed impacts of training data composition on model generalization. Despite IC_50_ measurements being generally considered less reliable due to their single-point assay nature, removing pIC_50_ data from training notably decreased predictive accuracy, particularly for pIC_50_ predictions alone (Pearson *r* reduced from 0.57 to 0.49; **Supplementary Table 3**). Given the substantial proportion of pIC_50_ data (approximately 33%) in the original training set (**Supplementary Table 1**), their exclusion likely reduced dataset diversity and consequently impacted the model’s generalizability. These results highlight the beneficial role of including diverse affinity types in enhancing DDMuffin’s predictive robustness.

To identify structural factors driving predictive accuracy, we applied t-distributed stochastic neighbour embedding (t-SNE) dimensionality reduction to computed descriptive features. Protein-ligand complex-derived features (**Supplementary Note 1**) formed clear clustering patterns corresponding to protein classes (**Fig. 2b**, left panel). Further analysis highlighted distinct residue and pocket polarity distributions among these classes (**Supplementary Fig. 2**). Lysine residues were enriched in lyases, likely reflecting their catalytic roles mediated through positively charged transition-state interactions. Chaperones exhibited an abundance of isoleucine residues, aligning with their reliance on hydrophobic core stability during protein folding. High glutamic acid content and pocket polarity observed in chaperones, hydrolases, transferases, and oxidoreductases indicate their shared preference for highly polar active-site environments critical for ligand binding. Histidine enrichment in metal-containing proteins aligns with its established role in metal ion coordination. Inclusion of ligand-centric descriptors provided limited enhancement in clustering clarity, although ligand classes remained distinguishable (**Fig. 2b**, right panel; **Supplementary Fig. 3)**.

When benchmarking against established predictive methods including AutoDock Vina^8^, Random Forest (RF)-Score^9^, InteractionGraphNet (IGN)^10^, and DeepDTA^11^, DDMuffin exhibited superior predictive performance on the LP-PDBBind test set, achieving the lowest Root Mean Squared Error (RMSE = 1.48) among all methods and a competitive Pearson correlation (*r* = 0.54), slightly trailing RF-Score (*r* = 0.58) (**Fig. 2c**). Detailed comparative metrics are available in **Supplementary Table 4**.

Permutation importance analyses on the LP-PDBBind^29^ test set further revealed distinct contributions from DDMuffin’s architecture components and descriptive features (**Fig. 2d**, **Methods**). The Descriptive Features block demonstrated the highest single-module impact (ΔPearson *r* = 0.23), followed by Graph-based Signatures (0.05), MoLFormer^40^ embeddings (0.03), and GCN-based modules (0.03). The combined permuting of Graph Block components (Graph-based Signatures and GCN, ΔPearson *r* = 0.11) and the integrated 1D Block (Descriptives and MoLFormer, ΔPearson *r* = 0.35) showed synergistic effects, with larger performance reductions than individual permutations combined. Simultaneous permutation of all modules produced the largest performance decrease (ΔPearson *r* = 0.49), underscoring significant interdependencies among architecture components. Additionally, combining ligand-protein complex descriptors with ligand 2D descriptors significantly improved predictive accuracy over individual ligand-only descriptors (**Fig. 2d**, lower panel; **Supplementary Table 5**; **Supplementary Note 1**).

Collectively, these findings validate DDMuffin’s ability to extract biologically meaningful interaction patterns from diverse protein-ligand datasets, providing comprehensive insights into feature-level contributions crucial for accurate predictive modelling.

### Benchmarking on external affinity test sets uncovers ligand binding determinants across protein conformational states

To assess the generalization capabilities of DDMuffin beyond the training and fine-tuning datasets, we conducted comprehensive evaluations using independent external test sets, namely BDB2020+^35^, EGFR^34^, and SARS-CoV-2 main protease (Mpro)^33^. These datasets were curated from the same source as LP-PDBBind^29^, to minimize overlap with the LP-PDBBind training data, ensuring a rigorous evaluation of model generalizability (**Method**). DDMuffin exhibited robust predictive performance across these datasets, achieving Pearson correlation coefficients (*r*) of 0.38, 0.56, and 0.69 for BDB2020+, EGFR, and Mpro, respectively (**Fig. 3a**). Notably, excluding the top 10% highest-residual predictions improved these correlations to 0.49, 0.67, and 0.80, highlighting the model’s capability to reliably predict typical binding affinities while isolating challenging cases.

**Fig. 3.**
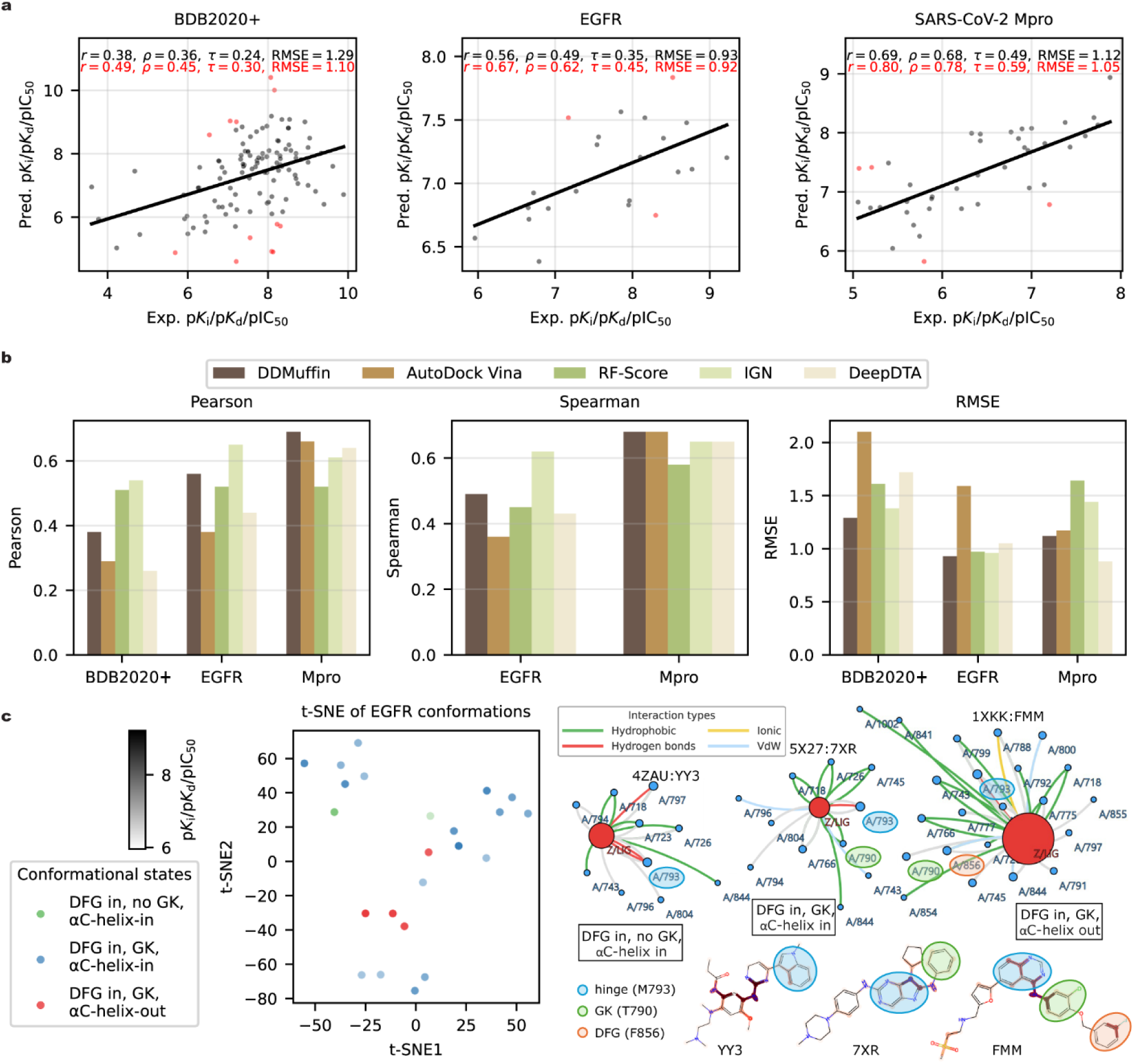
Performance evaluation and structural insights from DDMuffin on external affinity blind test sets. **a)** Scatter plots comparing predicted and experimental binding affinities (p*K*_i_, p*K*_d_, pIC_50_) for BDB2020+^35^, EGFR^34^, and SARS-Cov-2^33^ Mpro test sets. Performance metrics (Pearson (*r*), Spearman (*ρ*), Kendall (*τ*) correlations, and RMSE) are shown for complete datasets (black text) and after exclusion of the top 10% outliers (red dots and text). **b)** Comparative benchmarking of DDMuffin against established computational tools. DDMuffin was compared against AutoDock Vina^8^, Random Forest (RF)-Score^9^, InteractionGraphNet (IGN)^10^, and DeepDTA^11^ on BDB2020+, EGFR and Mpro (**Supplementary Table 7**). Performance of benchmarking methods were directly taken from the LP-PDBBind paper^29^. **c)** Structural analysis of EGFR conformational states. Left: t-SNE clustering identifies three EGFR conformational states distinguished by GK residue accessibility and αC-helix orientation, coloured accordingly and shaded by affinity strength. Right: Representative interaction networks for selected ligand-protein complexes (4ZAU:YY3, 5X27:7XR, 1XKK:FMM) highlighting specific residue interactions. Ligands are shown as red circles, protein residues as blue circles, and interactions are color-coded according to type: hydrophobic (green), ionic (yellow), hydrogen bonds (red), and van der Waals (VdW; light blue). Ligand molecular representations are highlighted based on atomic importance identified by GCN node weights.

Residual analysis revealed that prediction inaccuracies in the BDB2020+ dataset predominantly involved membrane protein complexes with ligands characterized by higher lipophilicity and molecular weight, typically interacting through atypical or allosteric binding sites. Although membrane proteins represented only 21% (25/115) of the dataset, they accounted for 83% (10/12) of outlier predictions. While membrane protein ligand outliers exhibited slightly higher molecular weights (411.3 ± 125.3 Da) and lipophilicity (log*P* 3.12 ± 2.09), and lower topological polar surface area (TPSA 76.2 ± 26.9 Å²) compared to non-outliers (380.6 ± 167.5 Da, log*P* 2.48 ± 1.78, TPSA 84.2 ± 23.8 Å²), these physicochemical differences did not reach statistical significance (**Supplementary Table 6**). Critically, atypical ligand binding modes of membrane proteins, particularly allosteric interactions observed in type A γ-aminobutyric acid (GABA_A_) receptor complexes, emerged as primary drivers of prediction inaccuracies, whereas ligands exhibiting canonical, orthosteric interactions with moderate physicochemical properties were accurately modelled. These observations suggest that the combination of ligand physicochemical extremes and unconventional binding interactions, rather than membrane localization alone, primarily drives prediction errors.

Comparative benchmarking against established affinity prediction methods further highlighted the competitive advantage of DDMuffin. When evaluated alongside AutoDock Vina^8^, RF-Score^9^, IGN^10^, and DeepDTA^11^, DDMuffin consistently outperformed AutoDock Vina and DeepDTA across all tested datasets, achieving lower RMSE and higher correlation coefficients (**Fig. 3b**, **Supplementary Table 7**). While RF-Score and IGN showed competitive correlation metrics for the BDB2020+ dataset, and IGN notably on the EGFR dataset, DDMuffin maintained superior overall performance by balancing RMSE and correlation across diverse protein targets, consistent with earlier findings from LP-PDBBind test set evaluations (**Fig. 2c**, **Supplementary Table 4**).

Structural analyses of EGFR complexes were conducted to identify conformational features underpinning DDMuffin’s predictions. EGFR, a critical therapeutic kinase known to adopt multiple conformations relevant to drug efficacy and resistance, served as a prototypical case study. Using t-SNE visualizations based on structural descriptors, three EGFR conformational states (as annotated in KLIFS^41^) defined by gatekeeper (GK) residue accessibility and αC-helix orientation were distinctly resolved (**Fig. 3c**, left).

Notably, the DFG-in, αC-helix-out conformation (*n* = 4) formed a relatively cohesive cluster, indicating structural consistency, whereas the DFG-in, αC-helix-in (GK accessible) conformation (*n* = 17) displayed broader structural diversity. The DFG-in, αC-helix-in (GK inaccessible) conformation (*n* = 2) was underrepresented, limiting robust conclusions.

Analysis of representative ligand-protein complexes (4ZAU:YY3, 5X27:7XR, and 1XKK:FMM) revealed distinct interaction patterns involving key residues such as hinge Met793, gatekeeper Thr790, and DFG motif residue Phe856 (**Fig. 3c**, right top; **Supplementary Fig. 4**). The αC-helix-out conformation (1XKK:FMM) exhibited the extensive and diverse interactions involving multiple discrete ligand fragments identified as critical via atom-level GCN weights, likely due to occupancy of extended binding sites. Conversely, αC-helix-in conformations exhibited fewer interactions, concentrating ligand importance into continuous structural fragments (**Fig. 3c**, right bottom). These structural insights emphasize the significance of capturing conformational diversity to enhance predictive accuracy in kinase-targeted drug discovery.

### Fine-tuning enhances mutation effect predictions on protein-ligand affinity

To improve accuracy in predicting single-point mutation effects on binding affinity (ΔΔ*G*), we fine-tuned the DDMuffin model using the smaller, mutation-specific Platinum^36^ dataset (*n* = 796), which is nearly ten times smaller than the general LP-PDBBind training set. Rigorous benchmarking across four stringent cross-validation scenarios revealed moderate predictive performance, reflecting the inherent challenges and variability associated with mutation-induced affinity predictions. Under 10-fold cross-validation designed to minimize redundancy by clustering protein-ligand complexes based on protein sequence and ligand chemical similarities (**Methods**), initial correlations were modest (Pearson *r* = 0.20, Spearman *ρ* = 0.17, Kendall *τ* = 0.11; RMSE = 1.78 kcal mol⁻¹), though excluding the top 10% largest-residual outliers provided slight improvements (r = 0.23, ρ = 0.19, τ = 0.12; RMSE = 1.44 kcal mol⁻¹). Similar modest performance was consistently observed under leave-one-UniProt-out, leave-one-ligand-out, and leave-one-complex-out validations (**Fig. 4a**), underscoring the difficulty in generalizing predictions across diverse chemical and structural contexts from limited data.

**Fig. 4.**
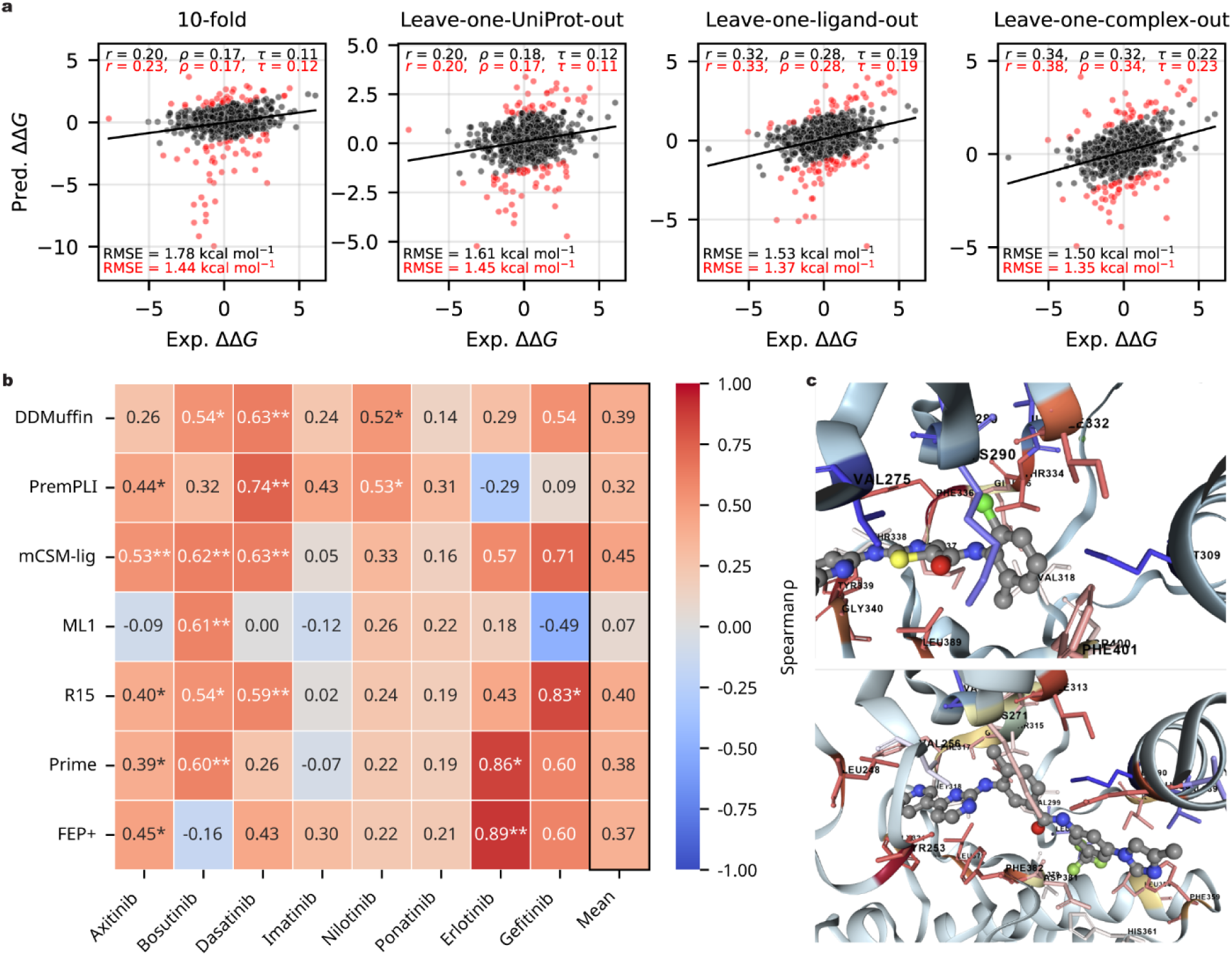
Prediction of affinity changes induced by single-point mutations. **a)** Scatter plots comparing predicted versus experimental binding affinity changes (ΔΔ*G*) on the Platinum^36^ dataset under four distinct cross-validation settings: 10-fold, leave-one-UniProt-out, leave-one-ligand-out, and leave-one-complex-out. Performance metrics include Pearson (*r*), Spearman (*ρ*), Kendall (*τ*) correlations, and root mean squared errors (RMSE). Metrics computed on the full dataset are shown in black, while metrics after excluding the top 10% largest residuals (marked as red dots) are highlighted in red. **b)** Heatmap benchmarking predictive performance (Spearman *ρ*) of DDMuffin against six established affinity prediction methods (PremPLI^19^, mCSM-lig^20^, ML1^25^, R15^26^, Prime^24^, and FEP+^24^) for eight different ABL1 kinase inhibitors in the TKI^24^ dataset. Predictions by benchmarking methods were taken from the PremPLI^19^ paper. Statistically significant correlations are indicated (\**P* value < 0.05, \*\**P* value < 0.01). Mean correlations across all inhibitors for each method are summarized in the rightmost column. **c)** Alanine scanning mapped onto ABL1 kinase domain structures bound to Type-I (Dasatinib, 4XEY, top) and Type-II inhibitors (Nilotinib, 3CS9, bottom). Key interacting residues are coloured based on ΔΔ*G* predictions by DDMuffin (red: reduced affinity, blue: enhanced affinity).

We further explored the impact of pre-training and fine-tuning strategies on DDMuffin’s predictive capabilities for mutation-induced affinity changes. The results indicated that fine-tuning on mutation-specific datasets primarily drives predictive performance, whereas the contribution of pre-training appeared relatively modest. Without initial pre-training on LP-PDBBind^29^, the correlations observed were only slightly lower (Pearson *r* = 0.18–0.31 across different validation settings) compared to scenarios involving pre-training (*r* = 0.20–0.34) (**Fig. 4a**, **Supplementary Fig. 5**). Additionally, direct calculation of ΔΔ*G* from separate wild-type and mutant affinity predictions using the pre-trained model yielded negligible correlation on the Platinum^36^ dataset and only a marginal correlation (*r* = 0.11) on the TKI dataset, underscoring the necessity of mutation-specific fine-tuning.

To maximize training data, we used HelixFold3^42^ to predict complex structures for ΔΔ*G* measurements from MdrDB^37^ lacking experimentally determined structures, incrementally improving predictive performance. Training exclusively on HelixFold3 predicted MdrDB structures produced modest correlations (*r* = 0.21) on the Platinum experimental dataset, which increased when incorporating HelixFold3 predicted TKI complexes (*r* = 0.26), and further improved when combined with experimental TKI complexes (*r* = 0.28). Interestingly, when tested on TKI dataset, relying exclusively on experimentally validated data in Platinum yielded stronger correlations (*r* = 0.38), while subsequent addition of HelixFold3 predictions from MdrDB slightly reduced correlation (*r* = 0.30). These results highlight the trade-off between data diversity and prediction accuracy, cautioning against indiscriminate integration of predicted complex structures.

DDMuffin’s predictive performance on mutation-driven affinity changes exhibited considerable variability across ligand and protein subsets under distinct validation approaches. For the two largest ligand subsets, flavin mononucleotide (FMN, *n* = 35) and guanosine diphosphate (GDP, *n* = 31), correlations under non-redundant 10-fold cross-validation were moderate (FMN *r* = 0.40; GDP *r* = 0.21). However, performance declined notably with stricter leave-one-ligand-out (FMN *r* = 0.29; GDP *r* = – 0.07) and leave-one-complex-out validations (FMN *r* = 0.04; GDP *r* = 0.12), accompanied by increased RMSEs (**Supplementary Fig. 6**). Analysis across protein subsets (*n* > 20, identified by UniProt IDs) also revealed variability. Leave-one-complex-out validation demonstrated strong correlations for several selected proteins (e.g., Aldo-keto reductase family 1 member B1 [P15121], *n* = 56, *r* = 0.75; Phenylethanolamine N-methyltransferase [P11086], *n* = 22, *r* = 0.71), but these correlations diminished significantly under other validations (**Supplementary Fig. 7-9**). These findings highlight the limited generalization capability from smaller mutation-specific datasets, emphasizing the need for more diverse data.

Comparative analyses underscored the competitive advantage of DDMuffin relative to existing computational methods. Across eight ABL1 kinase inhibitors, DDMuffin achieved consistently higher Spearman correlations (average Spearman *ρ* = 0.39), outperforming widely used methods such as PremPLI^19^ (average *ρ* = 0.32), Prime^24^ (average *ρ* = 0.38), and FEP+^24^ (average *ρ* = 0.37) (**Fig. 4b**).

Despite moderate correlations, DDMuffin consistently provided significant predictions for inhibitors such as Dasatinib (*P* value < 0.01), Bosutinib (*P* value < 0.01) and Nilotinib (*P* value < 0.05). Comparison on Pearson correlation and RMSE are detailed in **Supplementary Fig. 10**.

Alanine-scanning mutagenesis of the ABL1 active (Type I) and inactive (Type II) complexes reveals distinct resistance landscapes within the binding pockets (**Fig. 4c**). The type I (Dasatinib-bound, PDB ID: 4XEY) structure showed critical resistance hotspots (ΔΔ*G* ≤ −1 kcal mol⁻¹) around the hinge (F336) and neighbouring regions, including linker residues Y339–G340, gatekeeper T334, β5 residue I332 immediately N-terminal to the gatekeeper, and β7 residue L389 on the C-lobe; whereas mutations on αC (M309) and β2 (V275) modestly enhance affinity (ΔΔ*G* ≥ 0.5 kcal mol⁻¹) (**Supplementary Table 8**). Type II (Nilotinib-bound, PDB ID: 3CS9) hotspots (ΔΔ*G* ≤ −1 kcal mol⁻¹) shifted to the kinked Glycine-rich loop (G-loop) residue Y253, which interacts with the phosphate group of Adenosine triphosphate (ATP), β1 (L248), αC (E286), β5 (I313), and DFG motif (F382); mutations on αC (M290) modestly enhance affinity (ΔΔ*G* ≥ 0.5 kcal mol⁻¹) (**Supplementary Table 9**). These non-overlapping patterns underscore that Type I inhibitors depend primarily on hinge-gatekeeper contacts, whereas Type II inhibitors exploit the extended back pocket created in the αC-helix-out, DFG-out inactive conformation, illustrating distinct structural drivers of their resistance profiles.

### DDMuffin Web Server

The DDMuffin web server, accessible at https://biosig.lab.uq.edu.au/ddmuffin/, provides an intuitive and accessible platform for deep-learning based prediction of protein-ligand binding affinity and the impact of mutations (**Fig. 5**). Users begin by submitting a protein-ligand complex structure, either by uploading a PDB file or by specifying a PDB accession number, along with relevant ligand details (**Fig. 5a**).

**Fig. 5.**
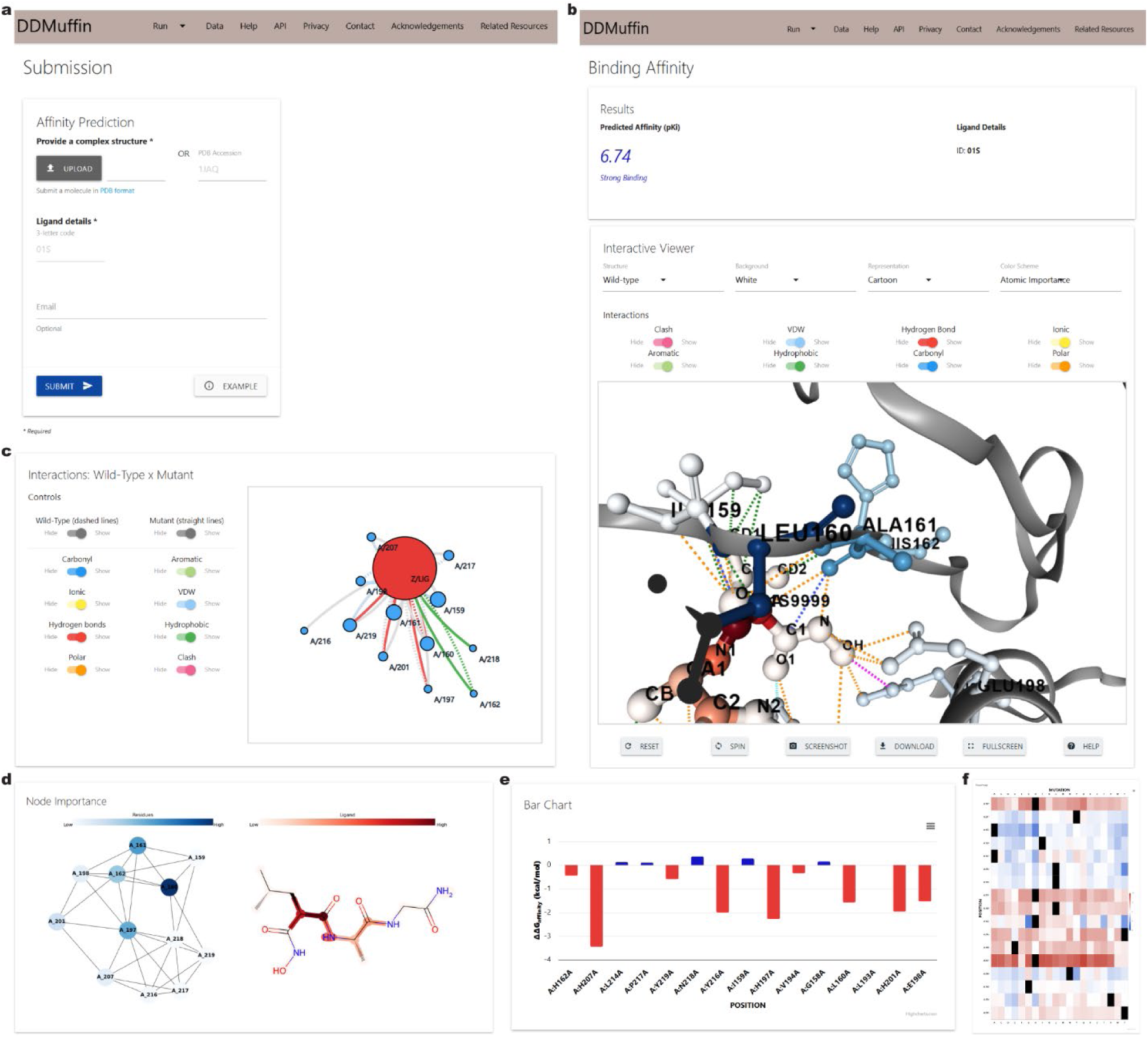
DDMuffin Web Server Interface. **a)** Submission interface for ‘Affinity Prediction’. Users upload protein-ligand complex structures in PDB format or specify a PDB accession number along with ligand details. **b)** ‘Affinity Prediction’ result page showing predicted affinity (p*K*_i_), ligand details, and an interactive molecular viewer that visualizes atomic interactions with customizable settings. **c)** Wild-type (dashed lines) and mutant (solid lines) interaction networks with selectable interaction types (carbonyl, ionic, hydrogen bonds, polar, aromatic, Van der Waals, hydrophobic, clash), shown on the ‘Single Mutation’ result page. **d)** Node importance visualization highlighting critical residues (left panel, blue) and ligand atoms (right panel, red) that significantly impact predicted binding affinity, color-coded as gradients. **e)** Bar chart summarizing ‘Alanine Scanning’ outcomes, indicating ΔΔ*G* upon alanine substitution at pocket residues (within 5Å of ligand atoms). Increased affinity (ΔΔ*G* > 0 kcal mol^-1^) is depicted in blue, while reduced affinity (ΔΔ*G* < 0 kcal mol^-1^) is shown in red. **f)** ‘Saturation Mutagenesis’ heatmap displaying predicted effects of all possible amino acid substitutions at each pocket residue on binding affinity, identifying critical residues influencing ligand binding affinity.

Following processing, results will be presented in a structured, multi-panel interface.

Upon submitting through the ‘Affinity Prediction’ tab, the result page displays the predicted binding affinity (p*K*_i_) and ligand information. A 3-dimensional interactive molecular viewer allows detailed exploration of the protein-ligand complex, featuring customizable visualization options such as atomic interactions, representation styles, and colour schemes indicating atomic importance (**Fig. 5b**).

Additionally, a dedicated 2-dimensional interaction network visualizes key contacts between ligand and protein residues, enabling selective viewing of specific interaction types, including hydrogen bonds, hydrophobic interactions, Van der Waals forces, and steric clashes (analogous to **Fig. 5c** but representing only the wild-type structure). Node importance visualizations further identify critical residues and ligand atoms through gradient colouring (blue for protein residues, red for ligand atoms) based on their contribution to binding affinity (**Fig. 5d**).

DDMuffin also supports detailed mutation analyses via ‘Single Mutation’ or ‘Mutation List’ submission panels through the ‘Mutation Effect’ tab. The ‘Single Mutation’ result page provides the predicted ΔΔ*G*, alongside mutation and ligand data. The interactive viewer available here is similar to the affinity result page but includes additional options for mutant structure visualization. Comparative interaction networks illustrate differences between wild-type and mutant interactions, clearly highlighting altered contacts (**Fig. 5c**). Node importance analyses are additionally provided for wild-type structures.

For comprehensive ‘Pocket Analysis’, users may perform ‘Alanine Scanning’ or ‘Saturation Mutagenesis’. ‘Alanine Scanning’ evaluates ΔΔ*G* resulting from alanine substitutions across all binding-pocket residues (within 5Å of ligand atoms), presented through bar charts summarizing residue-specific impacts (**Fig. 5e**). ‘Saturation Mutagenesis’ extends this analysis by systematically assessing affinity impacts of all possible amino acid substitutions at each pocket residue, visualized through informative heatmaps that efficiently highlight affinity-critical residues (**Fig. 5f**). Detailed usage instructions and further guidance are available on the DDMuffin help page: https://biosig.lab.uq.edu.au/ddmuffin/help.

## Discussion

We developed DDMuffin, a robust computational framework for predicting mutation-induced changes in protein-ligand binding affinity (ΔΔ*G*), achieving high accuracy across diverse validation datasets. Pre-trained on the extensive LP-PDBBind dataset, DDMuffin outperformed traditional and contemporary machine-learning methods, including AutoDock Vina^8^, RF-Score^9^, IGN^10^, and DeepDTA^11^ (**Fig. 2c**).

Crucially, DDMuffin effectively captured structural contributors of ligand binding, demonstrated by strong predictive performance on EGFR kinase inhibitors and SARS-CoV-2 main protease complexes (**Fig. 3a, b**), underscoring its broad applicability across therapeutic targets.

DDMuffin’s predictive accuracy largely stemmed from integrating advanced molecular representations: residue-level ProtT5^31^ embeddings, ligand pharmacophoric features, and protein-ligand interaction graphs derived from Arpeggio^32^. Permutation analyses revealed that each feature set and graph-based architecture significantly influenced performance, highlighting synergistic effects across architectural modules (**Fig. 2d, Supplementary Table 5**). These results emphasize the critical advantage of multi-scale features, combining local residue environments and detailed ligand interactions to provide essential predictive context.

An important consideration in interpreting DDMuffin’s affinity predictions is that the model was trained using diverse experimental affinity measurements, including pIC_50_, p*K*_d_ and p*K*_i_ values. Despite IC_50_ values being less reliable due to their single-point assay nature, their exclusion notably decreased predictive accuracy, particularly affecting p IC_50_ predictions (**Supplementary Table 3**). Given the substantial proportion of pIC50 data in the training set, their removal reduced dataset diversity, impacting model generalizability. Thus, the final prediction from DDMuffin should be interpreted as indicative of consensus affinity trends rather than exact binding constants.

Our analysis also identified specific challenges faced by DDMuffin, notably in predicting affinities for lyases, ligases, and membrane transport proteins (**Supplementary Table 2**). The disproportionate representation of these classes among outliers suggests that their unique binding mechanisms, such as cofactor-dependent catalysis and conformational plasticity, are inadequately captured by the current static atom-level descriptors. Moreover, prediction inaccuracies were prominent in membrane protein complexes exhibiting unconventional allosteric interactions, for example, the atypical binding modes in the GABA_A_ receptor (**Supplementary Table 6**). These observations highlight intrinsic limitations associated with static structural inputs and emphasize the importance of incorporating explicit modelling of dynamic or allosteric processes.

DDMuffin’s mutation-specific fine-tuning on the Platinum^36^ dataset provided moderate predictive performance, reflecting inherent challenges due to limited mutation-specific data. While fine-tuning markedly improved predictions compared to direct affinity subtraction methods, substantial variability persisted across ligand subsets and validation schemes. For example, predictive accuracy varied notably for flavin mononucleotide (FMN) and guanosine diphosphate (GDP) under different cross-validation settings (**Supplementary Fig. 6**). Additionally, protein-specific analyses revealed substantial variability across UniProt-defined subsets, indicating a critical need for larger and more diverse mutation-specific datasets (**Supplementary Fig. 7-9**).

The inclusion of computationally predicted structures from HelixFold3^42^ presented both benefits and limitations. Predicted structures improved model performance where experimental data was sparse but occasionally reduced accuracy compared to high-quality experimental structures. This finding emphasizes careful curation and validation of computationally derived complexes to balance dataset diversity with precision.

Conformational analyses, particularly of EGFR complexes, revealed valuable insights into structural aspects underlying binding predictions. Notably, distinct clustering of kinase conformational states (αC-helix-in and αC-helix-out) highlighted the importance of modelling conformational diversity for reliable predictions. Atom-level interaction analysis identified crucial residues and binding pockets that differentiated inhibitor binding patterns, providing mechanistic understanding that complements predictive accuracy (**Fig. 3c**). These insights are valuable for rational drug design, particularly in targeting specific kinase conformations.

### DDMuffin’s accessible and interpretable web interface (https://biosig.lab.uq.edu.au/ddmuffin/)

significantly enhances its practical utility, enabling detailed molecular visualizations, node importance analyses, and comprehensive mutation studies such as alanine scanning and saturation mutagenesis. Its intuitive design facilitates use by both specialists and non-specialists in drug discovery and personalized medicine, particularly addressing drug resistance and repurposing challenges. For example, systematically predicting resistance-conferring mutations can guide the design of resilient inhibitors, while broad affinity predictions can rapidly identify opportunities for drug repurposing.

Future developments in DDMuffin will benefit from incorporating dynamic structural ensembles and explicit modelling of allosteric effects, addressing current predictive limitations. Further enriching mutation-specific training data and systematically integrating experimentally validated structures will further enhance the model’s robustness and generalizability. Employing molecular dynamics simulations or enhanced sampling may also improve predictions by better capturing dynamic structural fluctuations and long-range allosteric effects. Ultimately, DDMuffin represents significant progress in computational drug discovery tools, combining predictive accuracy with interpretability to empower therapeutic decision-making against mutation-driven resistance and fostering innovative drug development.

## Methods

### Dataset preparation

During the pre-training phase, protein-ligand complexes in the LP-PDBBind^29^ Clean Level 1 subset were systematically curated from structures deposited in the RCSB Protein Data Bank, whereas additional validation sets (Mpro^33^, EGFR^34^, and BDB2020+^35^) were directly sourced and curated from the literature^29^. Specifically, BDB2020+ (*n* = 115) was compiled by matching high-quality binding affinity data from BindingDB^35^ with co-crystallized complexes deposited in the PDB after 2020, employing identical similarity control criteria used for LP-PDBBind development to minimize redundancy.

Similarly, EGFR (*n* = 23) and Mpro (*n* = 40) datasets were curated to exclude any complexes present in LP-PDBBind, However, SARS-CoV-1 protease entries, structurally similar to SARS-CoV-2 Mpro, were included. For fair comparison, benchmarking models including AutoDock Vina^8^, RF-Score^9^, IGN^10^, and DeepDTA^11^ were retrained on the same LP-PDBBind training set, and performance were directly taken from the paper^29^.

In the fine-tuning phase focusing specifically on mutation-induced affinity changes, experimental structures for complexes in the Platinum^36^ dataset and TKI-ABL1^24^ were similarly curated from their original sources. Due to the lack of experimentally resolved structures for the MdrDB^37^ dataset, these complexes were computationally predicted using HelixFold3^42^.

The ΔΔ*G* predicted by DDMuffin is defined as the difference in binding free energy between mutant and wild-type complexes, calculated as:

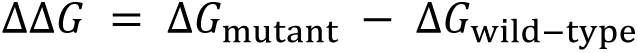

Positive ΔΔ*G* values indicate enhanced binding affinity (more favourable interactions), whereas negative values indicate reduced affinity (less favourable interactions). All ΔΔ*G* predictions from DDMuffin are expressed in kcal mol^-1^.

### Graph representation

Molecular graphs were generated using PyTorch Geometric^43^ to systematically encode structural and physicochemical interaction information. Three complementary graphs were constructed:

1. **Interaction Graph**: Nodes represent individual atoms annotated with 12 distinct pharmacophoric and functional types (e.g., hydrogen bond donors/acceptors, ionizable groups, hydrophobic groups, aromatic rings). Edges represent interatomic protein-ligand interactions annotated using Arpeggio^32^. Edge weights were assigned distinctively for 14 interaction types based on the relative energetic contribution required to disrupt each interaction (**Supplementary Table 10**).
2. **Protein Graph**: Nodes represent protein residues involved in binding interactions, annotated using high-dimensional ProtT5^31^ embeddings (1,024-dimensional) Edges connect residue pairs whose alpha carbons (Cα) are within 10 Å spatial proximity.
3. **Ligand Graph**: Nodes represent ligand atoms annotated by pharmacophoric types, with edges explicitly representing chemical bonds (single, double, triple, aromatic).

### Model architecture

DDMuffin employs a two-stage deep learning framework comprising pre-training (ProteinLigandGCN) and fine-tuning (ProtLigMutGCN) phases, leveraging GCNs with multiple feature extraction modules to predict mutation-induced ΔΔ*G*. The overall architecture is illustrated in **Supplementary Fig. 1**.

**Pre-training (ProteinLigandGCN).** To avoid over-training during the initial phase, the ProteinLigandGCN only uses four epochs, incorporating five primary modules:

Descriptive Features Module: Physicochemical and structural descriptors calculated via RDKit^44^ and binding-site features calculated via fpocket^45^ (detailed in **Supplementary Note 1**). Features are processed through fully connected (FC) layers with ReLU activation and dropout.

- MoLFormer^40^ Embedding Module: Molecular embeddings (768-dimensional) from MoLFormer, processed through FC layers with ReLU and dropout regularization.
- Graph-based Signature^20^ Module: Graph-derived pharmacophore features computed via a Cutoff Scanning Algorithm^46^. Atom pairs within distances ranging from 0.5 Å to 10 Å (incremental steps of 0.5 Å) were counted separately for eight pharmacophore categories (hydrophobic, aromatic, hydrogen bond donor, acceptor, positively charged, negatively charged, sulfur, and neutral), distinguishing between INTER (ligand-protein) and INTRA (ligand or protein) pairs. Resulting features are processed using convolutional layers (1D, kernel size = 2), adaptive average pooling, ReLU activation, and dropout.
- Graph Convolutional Network Module: Protein, ligand, and interaction graphs processed individually through GCNConv layers^43^ with sigmoid-activated edge weights, ReLU activations, and global mean pooling.
- Integration and Prediction Module: Outputs from the above modules concatenated into a unified embedding vector processed by a final FC layer predicting wild-type affinities.

**Fine-tuning (ProtLigMutGCN).** The fine-tuning stage was trained using 55 epochs, extending the ProteinLigandGCN by incorporating:

- Mutation Features Module: Mutation-specific descriptors (structural characteristics and amino acid indices^47,48^) computed using Biopython^49^ (**Supplementary Note 1**). Features processed through FC layers employing tanh and ReLU activation with dropout.
- Dual Prediction Strategy: Separate processing of wild-type and mutant features generates corresponding affinity predictions. Predicted p*K*_i_ values are first converted to inhibition constants (*K*_i_), which are subsequently used to calculate Gibbs free energies (Δ*G*) at physiological temperature (298.15 K), following the thermodynamic relationship:

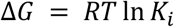

where *R* is the universal gas constant (1.987 × 10^-3^ kcal mol^-1^ K^-1^) and *T* is the absolute temperature in Kelvin. Mutation-induced ΔΔ*G* are then determined by taking the difference between the mutant and wild-type Gibbs free energies:

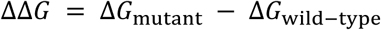

### Training data partitioning for hyperparameter optimization

To optimize hyperparameters via cross-validation, we divided the LP-PDBBind and Platinum training sets independently into ten non-redundant folds. To minimize redundancy and bias, we clustered protein-ligand complexes based on protein sequence and ligand chemical similarities. Protein sequence similarity was computed using MMseqs2^38^ with a threshold of 50% identity. Ligand similarity was quantified using Morgan fingerprints^39^ (radius = 2, 1024 bits) and Dice similarity^44^ scores, with a threshold of 0.95.

Additionally, an integrated overall similarity metric combining both protein and ligand similarity was calculated with a threshold of 0.6. We constructed a similarity graph, connecting complexes surpassing any similarity threshold, and identified connected components as clusters. These clusters were allocated into ten balanced folds without splitting, ensuring minimal overlap between training and validation subsets. Hyperparameter tuning scheme is detailed in **Supplementary Note 2**.

### Training and evaluation

Pre-training employed the LP-PDBBind dataset, followed by fine-tuning on the Platinum dataset. Model accuracy was evaluated using Pearson correlation (*r*), Spearman correlation (*ρ*), Kendall correlation (*τ*), and Root Mean Squared Error (RMSE) across independent datasets (TKI-ABL1^24^, MdrDB^37^, and BDB2020+^35^), along with additional validation datasets (SARS-CoV-2 Mpro^33^ and EGFR^34^).

### Permutation importance test

Permutation importance analyses were conducted to evaluate the individual contributions of different architectural blocks and descriptive feature sets to the overall predictive performance of DDMuffin on the LP-PDBBind test set. To achieve this, a baseline Pearson correlation was first established by evaluating the model on the original, unmodified test dataset.

Subsequently, permutation tests were performed separately for two groups: architectural components and descriptive feature sets. For the architectural components, key blocks (GCN, descriptive features, graph-based signatures, MoLFormer^40^ embeddings, as well as combined blocks) were systematically shuffled across the dataset. Similarly, for descriptive feature sets (**Supplementary Note 1**), groups such as ligand Lipinski and fragment descriptors (Lig1D), ligand graph and molecular surface area descriptors (Lig2D), and complex binding features were independently shuffled. Feature groups were permuted across samples while preserving intra-group correlations.

After each permutation, the model was evaluated on the modified dataset to determine the impact of disrupting the targeted block or feature group. The importance of each block or feature set was quantified by the change (Δ) in Pearson correlation relative to the baseline, thereby identifying the most influential contributors to DDMuffin’s predictive capability.

## Data availability

Data are available at https://biosig.lab.uq.edu.au/ddmuffin/datasets

## Acknowledgements

This work was supported by funding from the Australian Government Research Training Program Scholarship [to Y. Z.]; Investigator Grant from the National Health and Medical Research Council (NHMRC) of Australia [GNT1174405 to D.B.A.]; Victorian Government’s Operational Infrastructure Support Program. Funding for open access charge: NHMRC.

## Author contributions

Y.Z. conceived and led the study, designed and implemented the DDMuffin model, performed data analysis, built the web server, and wrote the manuscript. Y.M. supervised the project, provided critical insights, and assisted with the web server design. A.G.C.S. deployed and maintains the web server.

D.B.A. supervised the overall research, provided funding, and coordinated the study. All authors reviewed and approved the final manuscript.

## Ethics declarations

The authors declare no competing interests.

