## Supplementary Information for "Transfer learning across molecular graphs for predicting protein-ligand affinities and their changes upon mutations"

### Supplementary Tables

**Supplementary Table 1. Sample sizes by affinity measurement type across LP-PDBBind[1] datasets.**

| Affinity Measurement | Training | Validation | Testing |
|---|---|---|---|
| $IC_{50}$ | 2476 | 579 | 1020 |
| $K_i$ | 3023 | 962 | 1188 |
| $K_d$ | 2053 | 428 | 2078 |

**Supplementary Table 2. DDMuffin performance on LP-PDBBind[1] test set stratified by protein class.**

| Protein class* | Size | Pearson r | Spearman rho | Kendall tau | RMSE | Number of outliers | %Outlier by class | %All outliers |
|---|---|---|---|---|---|---|---|---|
| lyase | 429 | 0.284 | 0.258 | 0.172 | 2.270 | 147 | 34.27 | 34.27 |
| transport | 59 | 0.323 | 0.292 | 0.209 | 1.706 | 10 | 16.95 | 2.33 |
| membrane | 67 | 0.341 | 0.265 | 0.190 | 1.336 | 3 | 4.48 | 0.70 |
| transcription | 249 | 0.457 | 0.435 | 0.306 | 1.266 | 15 | 6.02 | 3.50 |
| other | 104 | 0.500 | 0.473 | 0.318 | 1.580 | 14 | 13.46 | 3.26 |
| ligase | 79 | 0.561 | 0.474 | 0.332 | 1.795 | 14 | 17.72 | 3.26 |
| transferase | 1703 | 0.600 | 0.575 | 0.404 | 1.300 | 94 | 5.52 | 21.91 |
| oxidoreductase | 177 | 0.633 | 0.644 | 0.454 | 1.150 | 6 | 3.39 | 1.40 |
| hydrolase | 1154 | 0.646 | 0.601 | 0.427 | 1.429 | 112 | 9.71 | 26.11 |
| chaperone | 183 | 0.665 | 0.552 | 0.395 | 1.284 | 8 | 4.37 | 1.86 |
| metal containing | 12 | 0.707 | 0.580 | 0.455 | 1.518 | 1 | 8.33 | 0.23 |
| isomerase | 54 | 0.707 | 0.724 | 0.519 | 1.514 | 3 | 5.56 | 0.70 |
| viral | 16 | 0.795 | 0.809 | 0.633 | 1.470 | 2 | 12.50 | 0.47 |

*Similar class distribution was observed in LP-PDBBind training and validation sets.

**Supplementary Table 3. DDMuffin predictive performance based on affinity measurement type.**

| **Training set** | **Test Set** | **Pearson *r*** | **RMSE** |
|---|---|---|---|
| $pIC_{50}$, $pK_d$ and $pK_i$ | $pK_d$ and $pK_i$ | 0.52 | 1.62 |
| $pIC_{50}$, $pK_d$ and $pK_i$ | $pIC_{50}$ | 0.57 | 1.31 |
| $pK_d$ and $pK_i$ | $pK_d$ and $pK_i$ | 0.49 | 1.66 |
| $pK_d$ and $pK_i$ | $pIC_{50}$ | 0.49 | 1.42 |

**Supplementary Table 4. Performance of protein-ligand affinity predictors on the LP-PDBBind[1] test set.**

| **Method*** | **Pearson *r*** | **RMSE** |
|---|---|---|
| DDMuffin | 0.54 | 1.48 |
| AutoDock Vina[2] | 0.45 | 2.56 |
| RF-Score[3] | 0.58 | 2.10 |
| IGN[4] | 0.51 | 2.16 |
| DeepDTA[5] | 0.49 | 2.29 |

*Performance other than DDMuffin was directly take from LP-PDBBind[1].

**Supplementary Table 5. Incremental contribution of feature sets to DDMuffin's accuracy on affinity prediction.** Feature descriptions are detailed in **Supplementary Note 1.**

| **Feature set** | **ΔPearson *r*** |
|---|---|
| Lig1D | 0.0176 |
| Complex | 0.0575 |
| Lig2D | 0.0803 |
| Lig1D + Complex | 0.1056 |
| Lig1D + Lig2D | 0.1396 |
| Lig2D + Complex | 0.1902 |
| All (Lig1D + Lig2D + Complex) | 0.2312 |

**Supplementary Table 6. Comparative features of membrane protein Outliers vs non-outliers in BDB2020+.**

| PDB ID | Ligand (Code) | MW (Da) | logP | TPSA (Å²) | Outlier | Binding Mode |
|---|---|---|---|---|---|---|
| 7QNE | Ro15-4513 (EIE) | 327.32 | 1.82 | 114.74 | Yes | Allosteric (benzodiazepine site on $GABA_A$) |
| 8DD2 | Zolpidem (R5R) | 342.45 | 1.21 | 37.61 | Yes | Allosteric (benzodiazepine site on $GABA_A$) |
| 8DD3 | DMCM (R63) | 332.41 | 2.93 | 73.44 | Yes | Allosteric (benzodiazepine site on $GABA_A$) |
| 7CKY | DRD1 agonist (G3U) | 349.40 | 1.74 | 86.21 | Yes | Orthosteric (GPCR active site) |
| 7DFL | Histamine (HSM) | 589.18 | 4.56 | 54.70 | Yes | Orthosteric (GPCR active site) |
| 7EJK | Oxymetazoline (J5C) | 211.33 | 1.58 | 48.91 | Yes | Orthosteric (GPCR active site) |
| 7EW2 | ISO-prostaglandin analog (J89) | 511.03 | 5.63 | 113.01 | Yes | Orthosteric (GPCR active site) |
| 7X2D | Tavapadon (86W) | 537.06 | 6.36 | 76.98 | Yes | Orthosteric (GPCR active site) |
| 6LR4 | ESF inhibitor (ESF) | 361.44 | 0.38 | 98.74 | Yes | Orthosteric (enzyme active site) |
| 7RJE | ZL5 antagonist (ZL5) | 551.36 | 4.97 | 57.48 | Yes | Orthosteric (GPCR active site) |
| **Outlier mean ± SD** | – | **411.3 ± 125.3** | **3.12 ± 2.09** | **76.2 ± 26.9** | – | – |
| 7CMU | Pramipexole (G6L) | 111.15 | -0.09 | 50.94 | No | Orthosteric (GPCR active site) |
| 7CMV | Labetalol analog (G6O) | 169.18 | 0.09 | 41.93 | No | Orthosteric (GPCR active site) |
| 7EJ0 | Norepinephrine (E5E) | 258.37 | 3.62 | 86.71 | No | Orthosteric (GPCR active site) |
| 7EJ8 | Brimonidine (J59) | 654.61 | 2.96 | 62.53 | No | Orthosteric (GPCR active site) |
| 7JVR | Bromocriptine (08Y) | 356.5 | 3.7 | 114.78 | No | Orthosteric (GPCR active site) |
| 7T6S | FPR2 antagonist (FUI) | 384.87 | 4.6 | 68.06 | No | Orthosteric (GPCR active site) |
| 7XTB | Serotonin (SRO) | 176.22 | 1.37 | 62.04 | No | Orthosteric (GPCR active site) |
| 7CX2 | Prostaglandin E2 (P2E) | 249.31 | 1.94 | 94.83 | No | Orthosteric (GPCR active site) |
| 7EW3 | Sphingosine-1-P (S1P) | 451.49 | 1.37 | 113.01 | No | Orthosteric (GPCR active site) |
| 7F58 | THIQ (1I8) | 415.19 | 3.26 | 92.15 | No | Orthosteric (GPCR active site) |
| 7T6B | Sphingosine-1-P (S1P) | 393.41 | 1.23 | 113.01 | No | Orthosteric (GPCR active site) |
| 7X2C | L-745,870 (G3C) | 665.77 | 4.42 | 72.72 | No | Orthosteric (GPCR active site) |
| 7Y5T | MRK-560 (IGD) | 517.93 | 4.67 | 80.31 | No | Orthosteric (enzyme active site) |
| 7XR6 | WAY-213613 (GJ0) | 385.5 | -0.3 | 101.65 | No | Orthosteric (transporter active site) |
| 7VSI | Empagliflozin (7R3) | 519.97 | 4.3 | 108.61 | No | Orthosteric (transporter active site) |
| **Non-outlier mean ± SD** | – | **380.6 ± 167.5** | **2.48 ± 1.78** | **84.2 ± 23.8** | – | – |

**Supplementary Table 7. Comparative benchmarking of DDMuffin and established affinity prediction methods.**

| **Dataset** | **Method*** | **Pearson *r*** | **Spearman *ρ*** | **RMSE** |
|---|---|---|---|---|
| BDB2020+ | DDMuffin | 0.38 | 0.36 | 1.29 |
| | AutoDock Vina | 0.29 | - | 2.10 |
| | RF-Score | 0.51 | - | 1.61 |
| | IGN | 0.54 | - | 1.38 |
| | DeepDTA | 0.26 | - | 1.72 |
| EGFR | DDMuffin | 0.56 | 0.49 | 0.93 |
| | AutoDock Vina | 0.38 | 0.36 | 1.59 |
| | RF-Score | 0.52 | 0.45 | 0.97 |
| | IGN | 0.65 | 0.62 | 0.96 |
| | DeepDTA | 0.44 | 0.43 | 1.05 |
| Mpro | DDMuffin | 0.69 | 0.68 | 1.12 |
| | AutoDock Vina | 0.66 | 0.68 | 1.17 |
| | RF-Score | 0.52 | 0.58 | 1.64 |
| | IGN | 0.61 | 0.65 | 1.44 |
| | DeepDTA | 0.64 | 0.65 | 0.88 |

*Performance other than DDMuffin was directly take from LP-PDBBind[1].

**Supplementary Table 8. ΔΔ*G* outcomes of alanine-scanning mutations predicted by DDMuffin on Dasatinib bound ABL1 active (Type I) complexes (PDB ID 4XEY).**

| **Wild-type** | **Residue number** | **Position*** | **Mutant** | **Ligand** | **Affinity prediction** | **ΔΔ*G* prediction (kcal mol$^{-1}$)** | **Change in affinity** |
|---|---|---|---|---|---|---|---|
| **PHE** | **336** | **Hinge** | **ALA** | **Dasatinib** | **6.76** | **-1.44** | **Decreasing** |
| **TYR** | **339** | **Linker** | **ALA** | **Dasatinib** | **6.76** | **-1.33** | **Decreasing** |
| **THR** | **334** | **GK** | **ALA** | **Dasatinib** | **6.76** | **-1.18** | **Decreasing** |
| **LEU** | **389** | **β7** | **ALA** | **Dasatinib** | **6.76** | **-1.11** | **Decreasing** |
| **ILE** | **332** | **β5** | **ALA** | **Dasatinib** | **6.76** | **-1.07** | **Decreasing** |
| **GLY** | **340** | **Linker** | **ALA** | **Dasatinib** | **6.76** | **-1.0** | **Decreasing** |
| LEU | 267 | β1 | ALA | Dasatinib | 6.76 | -0.82 | Decreasing |
| PHE | 401 | DFG | ALA | Dasatinib | 6.76 | -0.67 | Decreasing |
| MET | 337 | Hinge | ALA | Dasatinib | 6.76 | -0.52 | Decreasing |
| GLU | 335 | Hinge | ALA | Dasatinib | 6.76 | -0.47 | Decreasing |
| GLY | 268 | G-loop | ALA | Dasatinib | 6.76 | -0.32 | Decreasing |
| ASP | 400 | DFG | ALA | Dasatinib | 6.76 | -0.26 | Decreasing |
| VAL | 318 | B-loop | ALA | Dasatinib | 6.76 | -0.11 | Decreasing |
| THR | 338 | Linker | ALA | Dasatinib | 6.76 | -0.08 | Decreasing |
| LYS | 290 | β3 | ALA | Dasatinib | 6.76 | 0.15 | Increasing |
| ILE | 333 | β5 | ALA | Dasatinib | 6.76 | 0.31 | Increasing |
| VAL | 289 | β3 | ALA | Dasatinib | 6.76 | 0.4 | Increasing |
| **VAL** | **275** | **β2** | **ALA** | **Dasatinib** | **6.76** | **0.54** | **Increasing** |
| **MET** | **309** | **αC** | **ALA** | **Dasatinib** | **6.76** | **0.57** | **Increasing** |

*Position annotations are obtained from KLIFS[6].

**Supplementary Table 9. ΔΔ*G* outcomes of alanine-scanning mutations predicted by DDMuffin on Nilotinib bound ABL1 inactive (Type II) complexes (PDB ID 3CS9).**

| Wild-type | Residue number | Position* | Mutant | Ligand | Affinity prediction | ΔΔ*G* prediction (kcal $mol^{-1}$) | Change in affinity |
|---|---|---|---|---|---|---|---|
| **TYR** | **253** | **G-loop** | **ALA** | **Nilotinib** | **7.68** | **-1.54** | **Decreasing** |
| **ILE** | **313** | **β5** | **ALA** | **Nilotinib** | **7.68** | **-1.24** | **Decreasing** |
| **LEU** | **248** | **β1** | **ALA** | **Nilotinib** | **7.68** | **-1.19** | **Decreasing** |
| **GLU** | **286** | **αC** | **ALA** | **Nilotinib** | **7.68** | **-1.17** | **Decreasing** |
| **PHE** | **382** | **DFG** | **ALA** | **Nilotinib** | **7.68** | **-1.09** | **Decreasing** |
| LEU | 370 | β7 | ALA | Nilotinib | 7.68 | -0.99 | Decreasing |
| PHE | 359 | β6 | ALA | Nilotinib | 7.68 | -0.83 | Decreasing |
| GLY | 321 | Linker | ALA | Nilotinib | 7.68 | -0.82 | Decreasing |
| ILE | 293 | B-loop | ALA | Nilotinib | 7.68 | -0.69 | Decreasing |
| GLU | 316 | Hinge | ALA | Nilotinib | 7.68 | -0.59 | Decreasing |
| PHE | 317 | Hinge | ALA | Nilotinib | 7.68 | -0.56 | Decreasing |
| LEU | 354 | αE | ALA | Nilotinib | 7.68 | -0.56 | Decreasing |
| ASP | 381 | DFG | ALA | Nilotinib | 7.68 | -0.43 | Decreasing |
| ILE | 314 | β5 | ALA | Nilotinib | 7.68 | -0.42 | Decreasing |
| LYS | 271 | β3 | ALA | Nilotinib | 7.68 | -0.37 | Decreasing |
| THR | 315 | GK | ALA | Nilotinib | 7.68 | -0.24 | Decreasing |
| VAL | 299 | B-loop | ALA | Nilotinib | 7.68 | -0.13 | Decreasing |
| MET | 318 | Hinge | ALA | Nilotinib | 7.68 | -0.11 | Decreasing |
| HIS | 361 | C-loop | ALA | Nilotinib | 7.68 | -0.07 | Decreasing |
| VAL | 379 | β8 | ALA | Nilotinib | 7.68 | -0.05 | Decreasing |
| VAL | 256 | β2 | ALA | Nilotinib | 7.68 | 0.02 | Increasing |
| LYS | 285 | αC | ALA | Nilotinib | 7.68 | 0.08 | Increasing |
| LEU | 298 | B-loop | ALA | Nilotinib | 7.68 | 0.08 | Increasing |
| VAL | 289 | αC | ALA | Nilotinib | 7.68 | 0.17 | Increasing |
| VAL | 270 | β3 | ALA | Nilotinib | 7.68 | 0.34 | Increasing |
| **MET** | **290** | **αC** | **ALA** | **Nilotinib** | **7.68** | **0.87** | **Increasing** |

*Position annotations are obtained from KLIFS[6].

**Supplementary Table 10. Interaction types and corresponding edge weights.** Protein-ligand interaction types identified by Arpeggio[7], and their corresponding edge weights used in the interaction graph representation. Edge weights represent the relative energetic contributions required to disrupt each interaction type.

| **Interaction type** | **Edge weight** |
|---|---|
| Covalent bond | 1.000 |
| Ionic interaction | 0.941 |
| Metal complex | 0.665 |
| Hydrogen bond | 0.540 |
| Halogen bond | 0.524 |
| Polar interaction | 0.514 |
| Weak hydrogen bond | 0.508 |
| Aromatic interaction | 0.508 |
| Hydrophobic interaction | 0.508 |
| Weak polar interaction | 0.497 |
| van der Waals interaction | 0.497 |
| Carbonyl interaction | 0.503 |
| van der Waals clash | 0.343 |
| Clash | 0.100 |

### Supplementary Figures

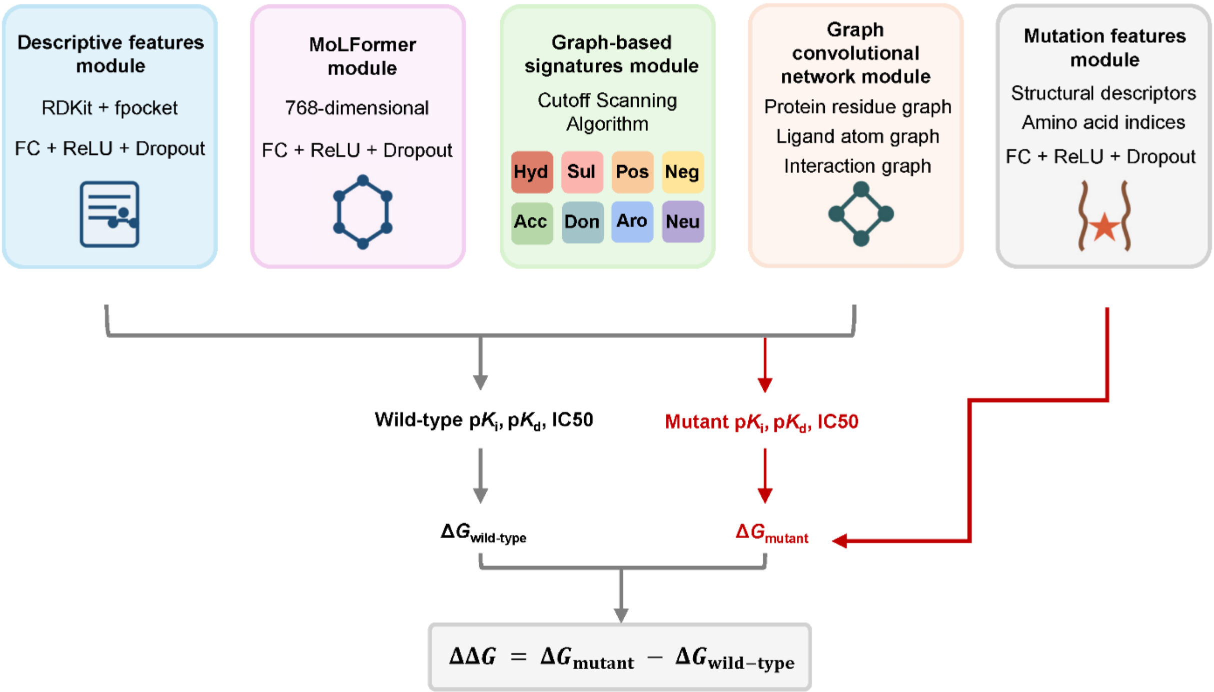

**Supplementary Fig. 1. Overview of the DDMuffin model architecture.** The DDMuffin approach utilizes a two-stage deep learning framework comprising pre-training and fine-tuning phases to predict mutation-induced changes in protein-ligand binding affinities ($\Delta\Delta G$). During pre-training, wild-type features are derived from physicochemical descriptors (RDKit[8], fpocket[9]), molecular embeddings (MoLFormer[10]), graph-based pharmacophore signatures (cutoff scanning algorithm[11,12]), and graph convolutional networks (protein, ligand, and interaction graphs). The fine-tuning phase incorporates mutation-specific structural descriptors and amino acid indices. Separate wild-type and mutant affinity predictions (p$K_i$, p$K_d$, $IC_{50}$) are converted into Gibbs free energies ($\Delta G$) to compute the final predicted affinity change ($\Delta\Delta G$).

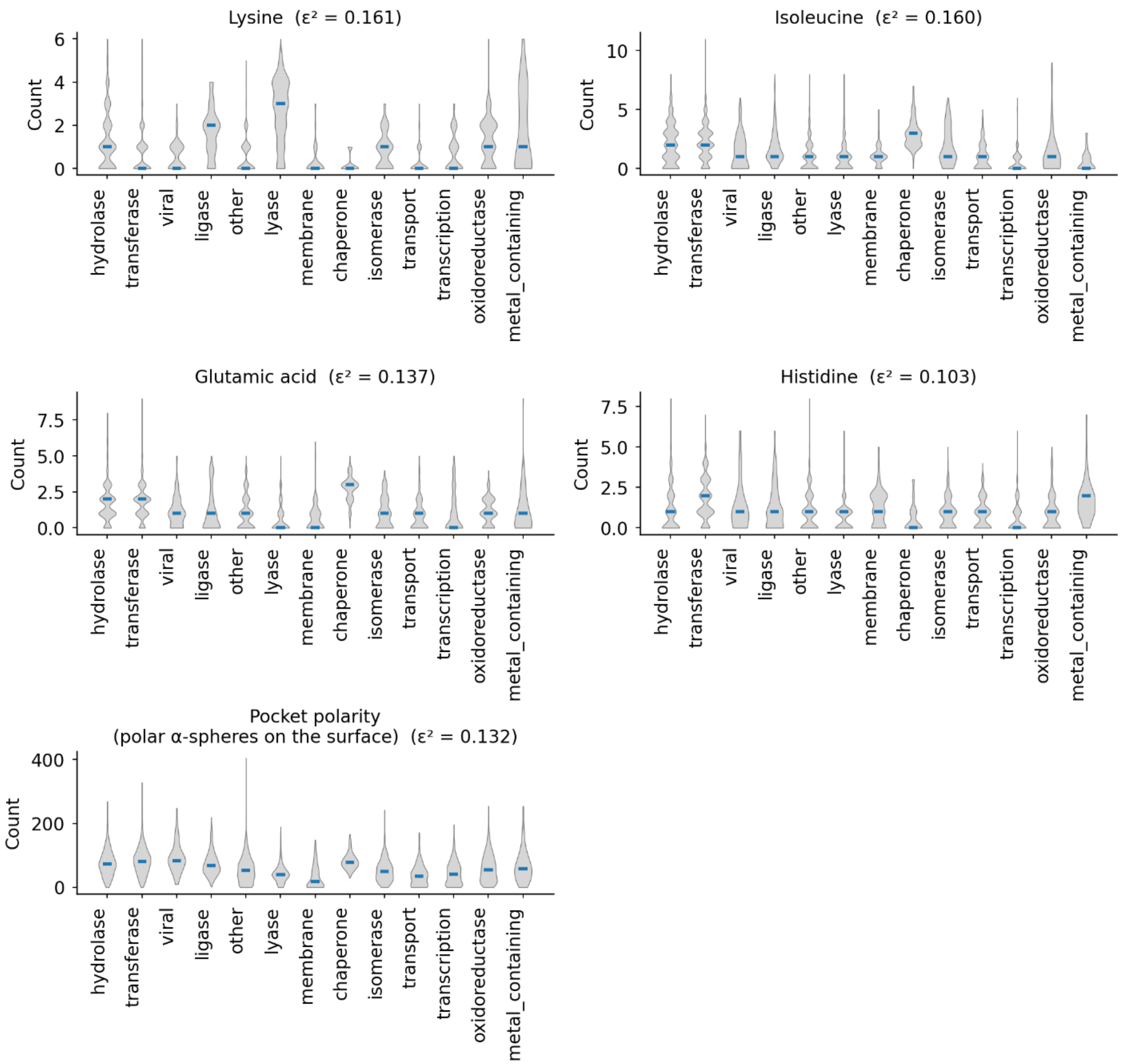

**Supplementary Fig. 2. Residue-specific and pocket-polarity signatures across protein functional classes within the LP-PDBBind dataset.** Violin plots show the distribution of lysine, isoleucine, glutamic-acid and histidine counts, plus binding-pocket polarity (number of polar α-spheres on the pocket surface), in 14 protein classes within the LP-PDBBind[1] dataset. Central blue bars mark the median. Kruskal-Wallis $\varepsilon^2$ values (in brackets) quantify the effect size of class-dependent differences. Values ≥ 0.14 are usually taken as a large effect (i.e., clear separation between classes), whereas 0.06–0.14 is moderate and < 0.06 is negligible.

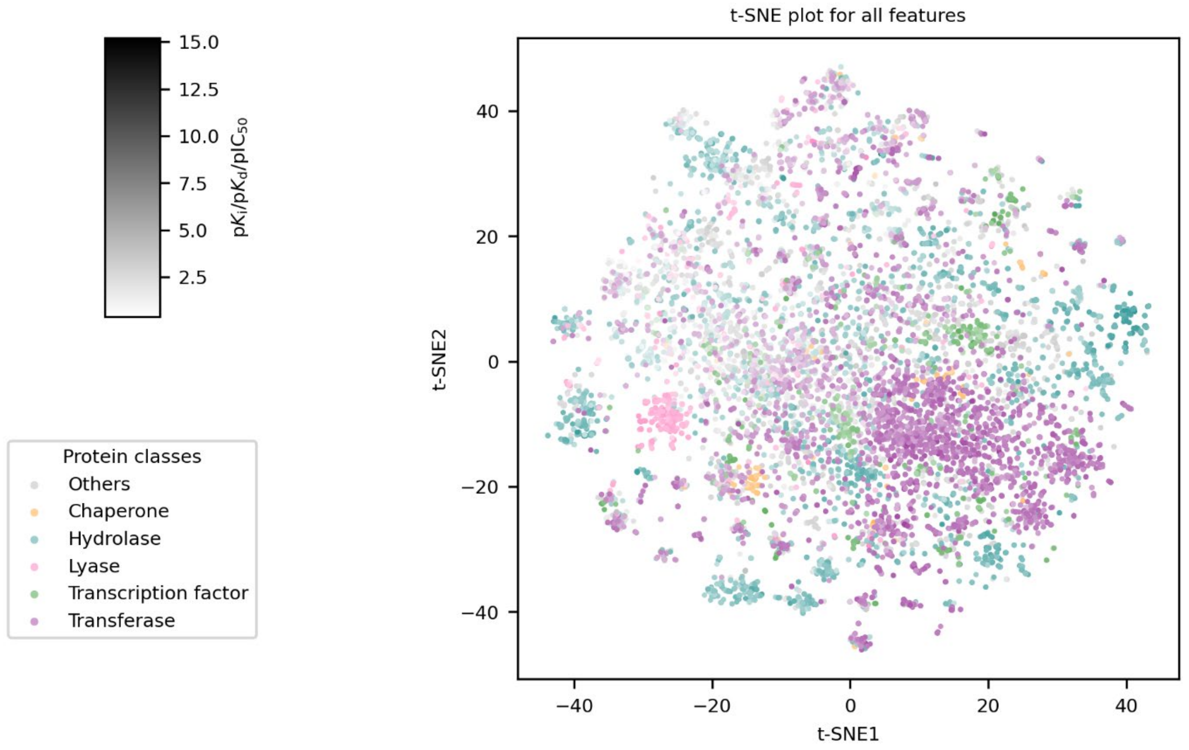

**Supplementary Fig. 3. t-SNE visualization of all descriptive features computed for the full LP-PDBBind dataset.** Points coloured according to protein classes (chaperone, hydrolase, lyase, transcription factor, transferase, and others), and shaded by affinity strength.

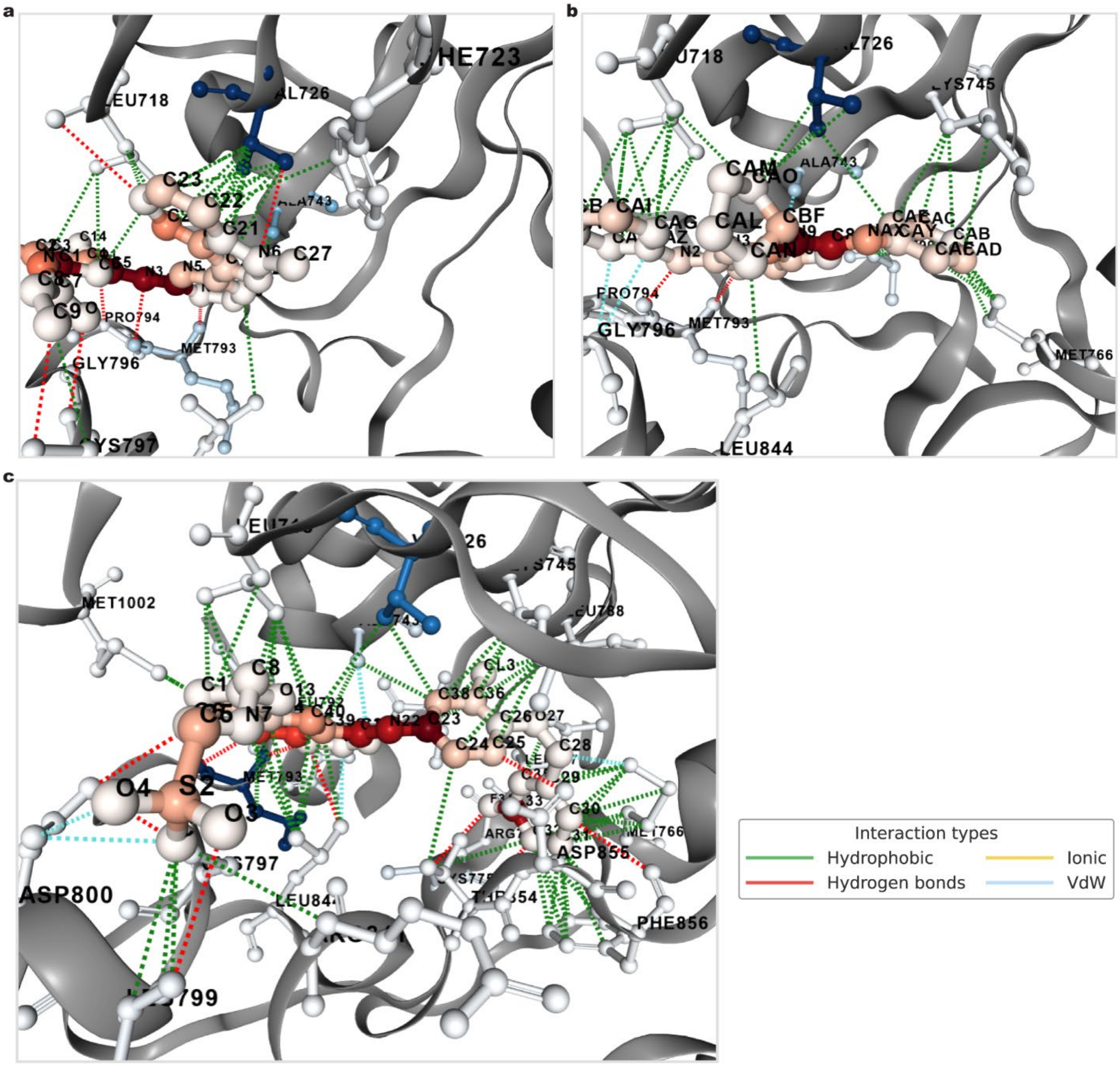

**Supplementary Fig. 4. Detailed three-dimensional representation of ligand-protein interactions in EGFR complexes corresponding to distinct conformational states.** **(a)** 4ZAU (ligand YY3, DFG-in, αC-helix-in, no gatekeeper accessibility), **(b)** 5X27 (ligand 7XR, DFG-in, αC-helix-in, gatekeeper accessible), and **(c)** 1XKK (ligand FMM, DFG-in, αC-helix-out, gatekeeper accessible). Residues involved in interactions are labelled explicitly. Interaction types are differentiated as hydrophobic (green), hydrogen bonds (red), ionic (yellow), and van der Waals (blue). Ligand atoms and protein residues are coloured according to GCN-derived atom node weights, emphasizing fragment contributions to binding affinity predictions.

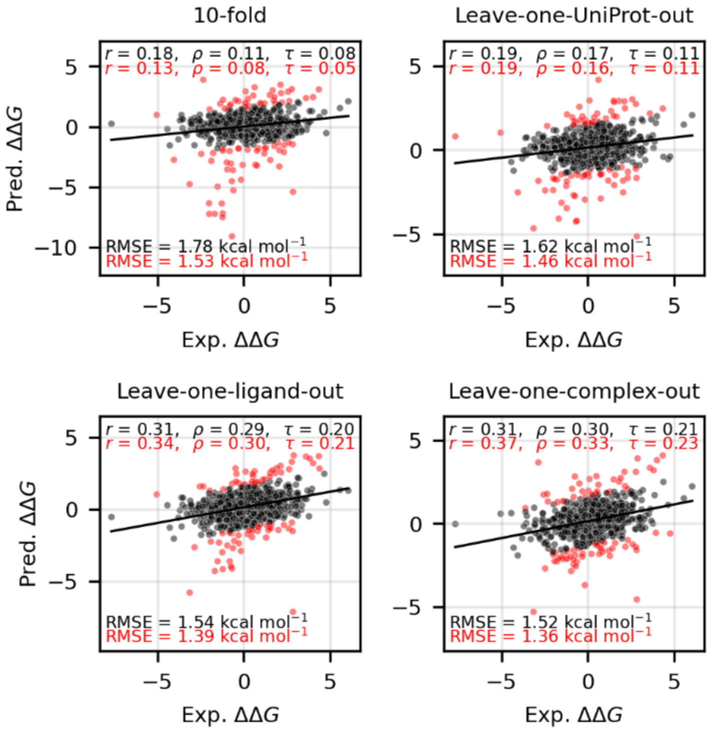

**Supplementary Fig. 5. Scatter plots comparing predicted versus experimental binding affinity changes (ΔΔ*G*) on the Platinum dataset without pre-training.** Performances were under four distinct cross-validation settings: 10-fold with minimized redundancy through protein sequence and ligand chemical similarity clustering, leave-one-UniProt-out, leave-one-ligand-out, and leave-one-complex-out. Performance metrics include Pearson ($r$), Spearman ($\rho$), Kendall ($\tau$) correlations, and root mean squared errors (RMSE). Metrics computed on the full dataset are shown in black, while metrics after excluding the top 10% largest residuals (marked as red dots) are highlighted in red.

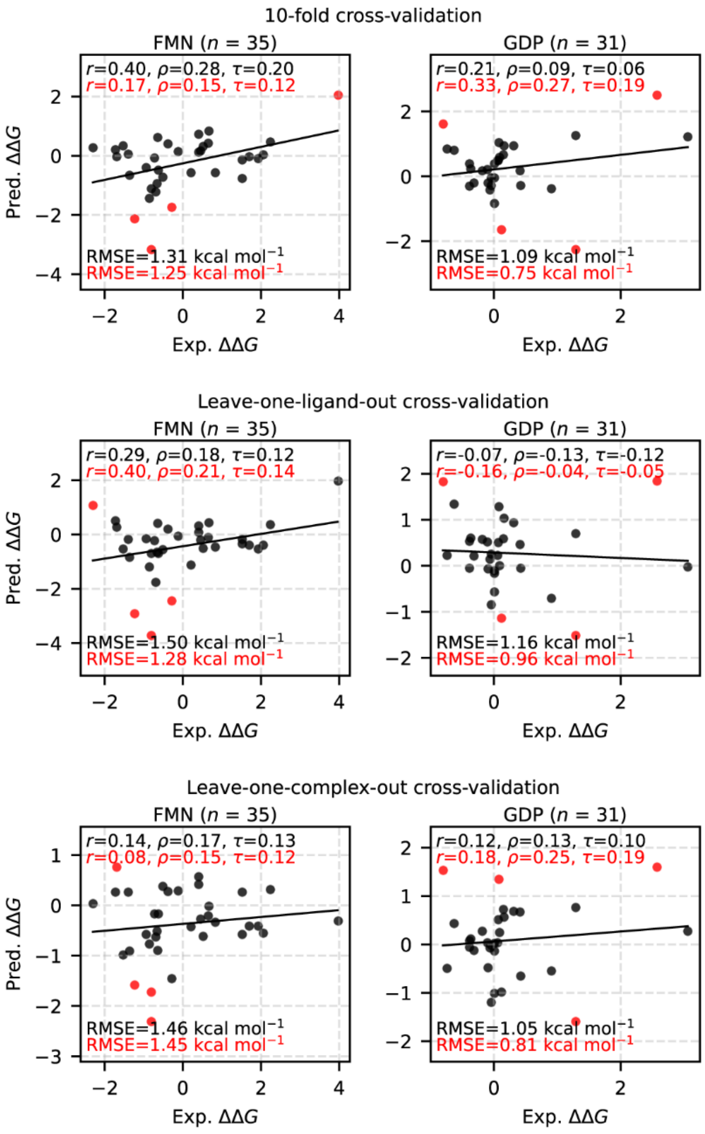

**Supplementary Fig. 6. Validation performance of DDMuffin on mutation-driven affinity prediction for the two largest ligand subsets (FMN, n = 35; GDP, n = 31).** Scatter plots depict predicted versus experimental mutation-induced binding affinity changes (ΔΔG) using three validation strategies: 10-fold cross-validation with minimized redundancy through protein sequence and ligand chemical similarity clustering (top row), leave-one-ligand-out cross-validation (middle row), and leave-one-complex-out cross-validation (bottom row). Performance metrics include Pearson (*r*), Spearman (*ρ*), Kendall (*τ*) correlations, and RMSE, calculated for the full datasets (black) and after excluding the top 10% largest-residual predictions (red points and annotations).

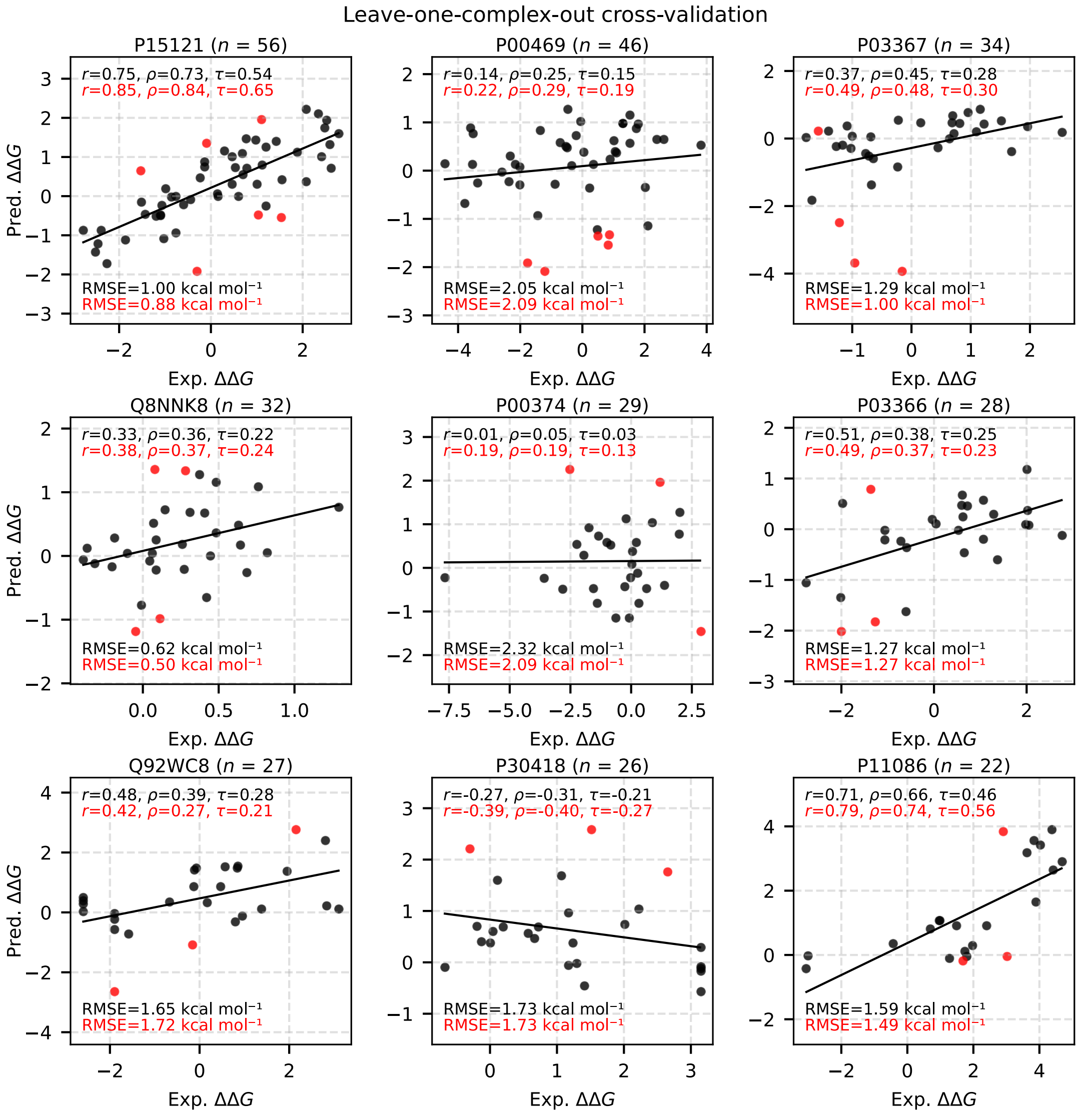

**Supplementary Fig. 7. Validation performance of DDMuffin on mutation-driven affinity prediction for protein subsets (leave-one-complex-out cross-validation).** Scatter plots illustrate predicted versus experimental mutation-induced affinity changes ($\Delta\Delta G$) for proteins with over 20 associated data points, identified by UniProt IDs: Aldo-keto reductase family 1 member B1 (AKR1B1, P15121, $n = 56$) from *Homo sapiens*; Thymidylate synthase (P00469, $n = 46$) from *Lacticaseibacillus casei*; Gag-Pol polyprotein (P03367, n = 34) of HIV-1; *pimB* protein (Q8NNK8, $n = 32$) from *Corynebacterium glutamicum*; Dihydrofolate reductase (P00374, $n = 29$) from *Homo sapiens*; Gag-Pol polyprotein (P03366, $n = 28$) of HIV-1, Cystine-binding periplasmic protein (Q92WC8, $n = 27$) from *Rhizobium meliloti*, Glycylpeptide N-tetradecanoyltransferase (P30418, $n = 26$) from *Candida albicans*, and Phenylethanolamine N-methyltransferase (P11086, $n = 22$) from *Homo sapiens*. Performance metrics shown are Pearson ($r$), Spearman ($\rho$), Kendall ($\tau$) correlations, and RMSE, calculated for full datasets (black) and after excluding the top 10% largest-residual predictions (red points and annotations).

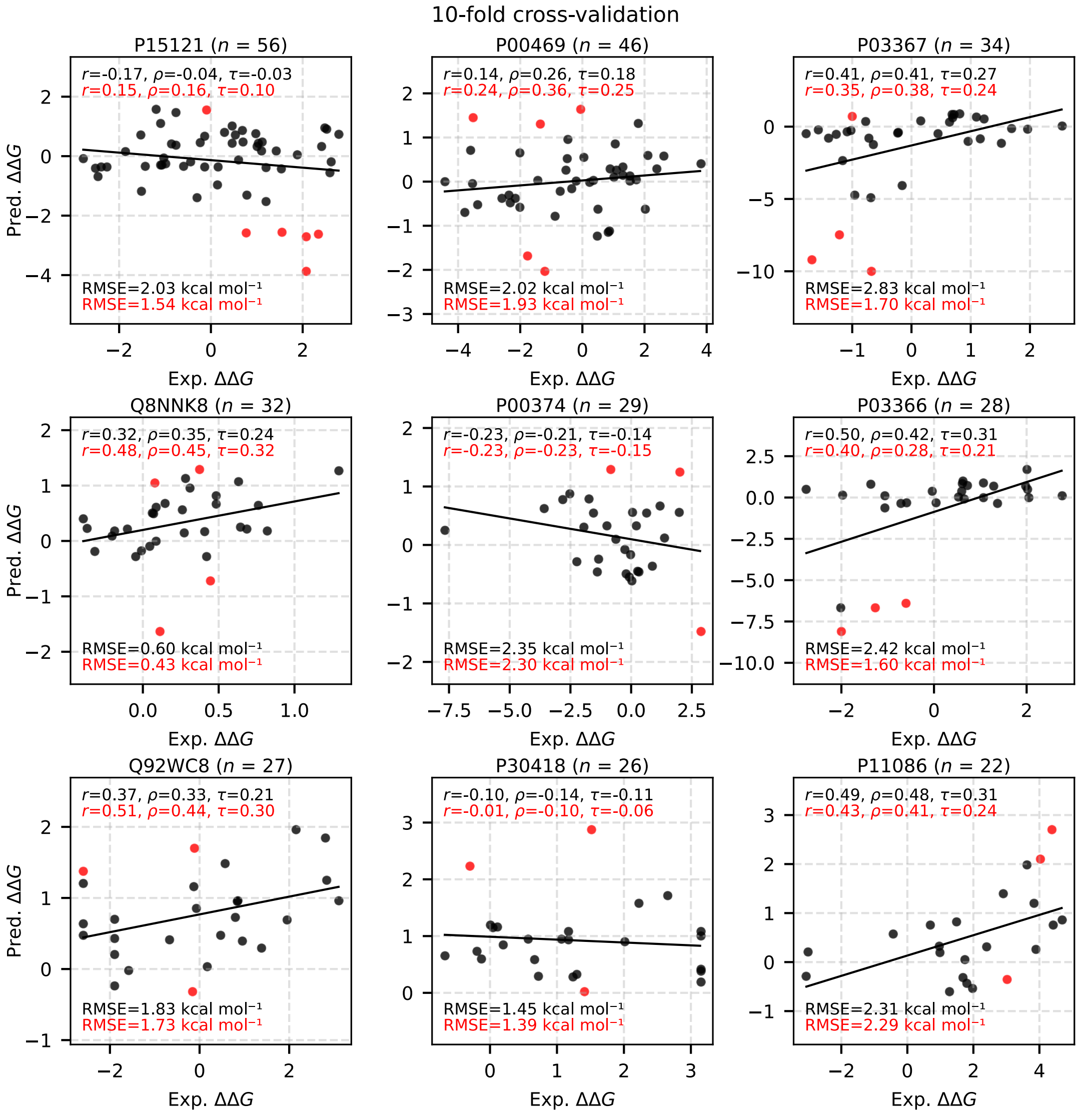

**Supplementary Fig. 8. Validation performance of DDMuffin on mutation-driven affinity prediction for protein subsets (10-fold cross-validation).** Scatter plots depict predictive accuracy for the same protein subsets as in **Supplementary Fig. 6**, using 10-fold cross-validation designed to minimize redundancy based on clustering by protein sequence and ligand chemical similarity.

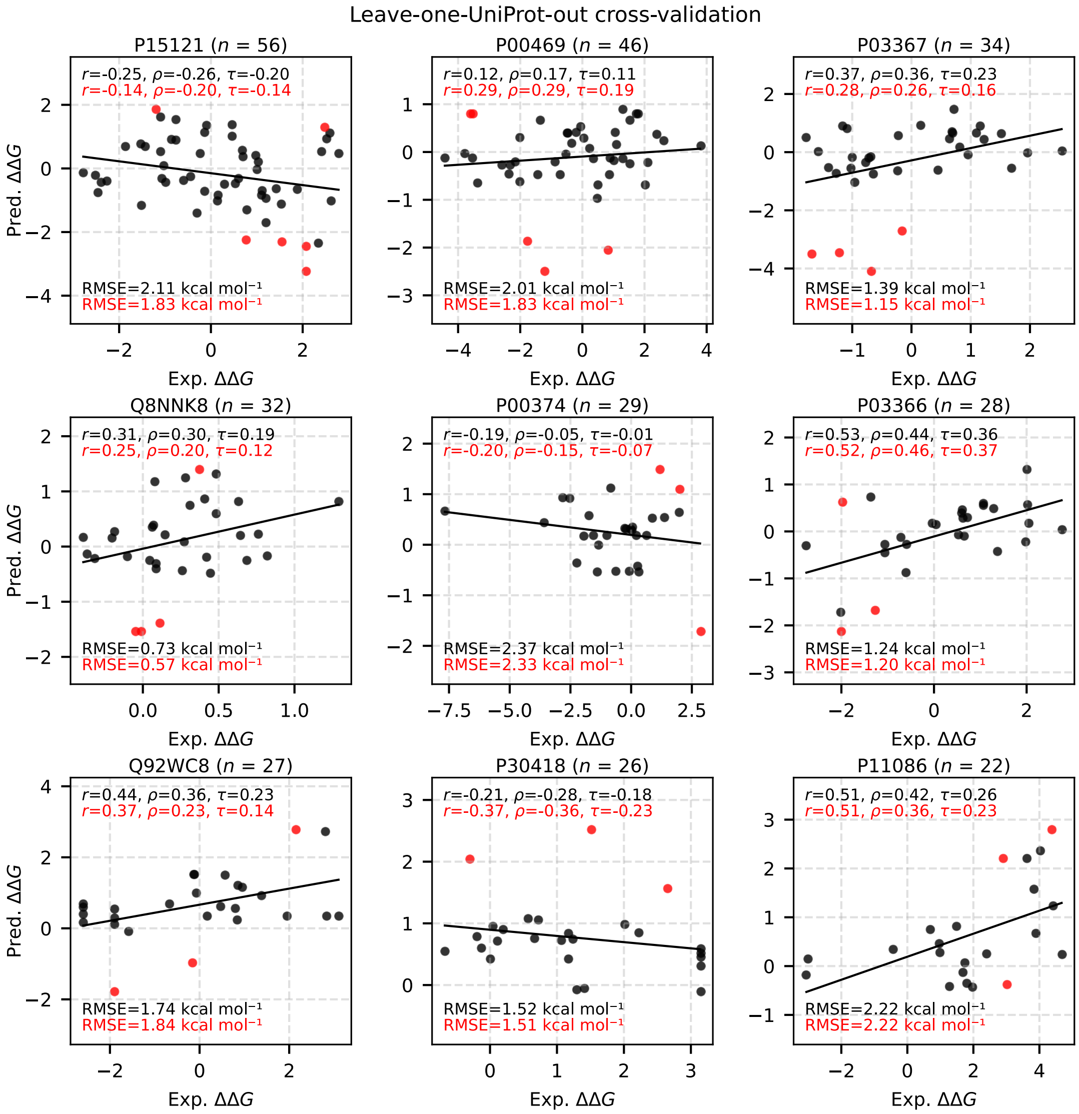

**Supplementary Fig. 9. Validation performance of DDMuffin on mutation-driven affinity prediction for protein subsets (leave-one-UniProt-out cross-validation).** Scatter plots illustrate predictive performance when entire protein subsets are held out from training, representing the strictest generalization test among the three validation strategies. Protein subsets are consistent with those shown in **Supplementary Fig. 6, 7**.

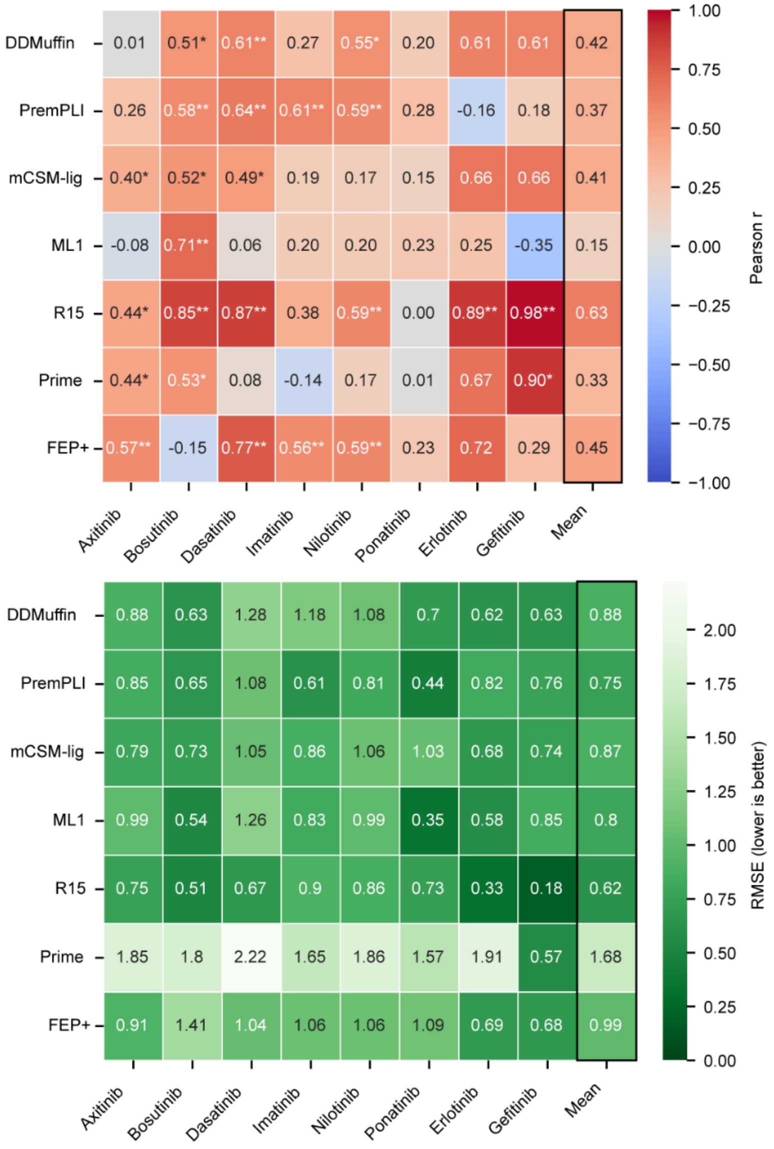

**Supplementary Fig. 10. Heatmap benchmarking predictive performance (Pearson r and RMSE) of DDMuffin against six established affinity prediction methods for eight different ABL1 kinase inhibitors in the TKI dataset.** Statistically significant correlations are indicated (**P* value < 0.05, ***P* value < 0.01). Mean correlations and RMSE across all inhibitors for each method are summarized in the rightmost columns. Predictions by benchmarking methods were taken from the PremPLI[13] paper.

### Supplementary Notes

#### Supplementary Note 1. Feature descriptions for DDMuffin

Detailed descriptions of features employed in the DDMuffin method for predicting protein-ligand interaction affinities and mutation-induced changes in binding affinities ($\Delta\Delta G$). The features are organized according to their use in the DDMuffin two-stage deep learning architecture (ProteinLigandGCN for pre-training and ProtLigMutGCN for fine-tuning).

Descriptive Features Module in ProteinLigandGCN:

1. Lig1D Features calculated via RDKit[8]:
   - Physicochemical descriptors: Molecular weight (MolWt), molecular log*P* (MolLog*P*), topological polar surface area (TPSA), rotatable bond count, heavy atom count, heteroatom count, hydroxyl groups (NHOHCount), nitrogen-oxygen groups (NOCount).
   - Pharmacophoric and toxicophoric fingerprints: Counts of specific functional groups (e.g., hydrogen bond donors/acceptors, aromatic rings, toxic groups).
2. Lig2D Features calculated via RDKit[8]:
   - Molecular complexity indices: Bertz complexity (BertzCT), Balaban's J index.
   - Connectivity and shape descriptors: Chi indices (Chi0, Chi0n, Chi0v, Chi1, Chi1n, Chi1v, Chi2n, Chi2v, Chi3n, Chi3v, Chi4n, Chi4v), Kappa shape indices (Kappa1, Kappa2, Kappa3), Labute accessible surface area (LabuteASA).
   - Partial atomic charge and surface area descriptors: PEOE_VSA, SMR_VSA, SlogP_VSA, and VSA EState indices.
3. Complex-based Features:
   - Interaction descriptors calculated via Arpeggio[7]: Non-covalent interactions including hydrogen bonds, halogen bonds, ionic interactions, hydrophobic contacts, aromatic and polar interactions, pi interactions, van der Waals interactions, and metal coordination.
   - Interaction graph-based descriptors calculated via iGraph[14]: Degree, betweenness, closeness, page rank, clustering coefficient, eccentricity, diameter, radius, centrality measures, entropy, and energy.
   - Binding site features calculated via fpocket[9]: Pocket volume, druggability scores, hydrophobicity, polarity, charge scores, surface metrics (asphericity, apolar/aspheric proportion, mean radius, solvent accessibility, local hydrophobic density), and amino acid composition within binding pockets.

Mutation Features Module in ProtLigMutGCN:

1. Structural features and environmental features of the mutated protein residues extracted from PDB structures, calculated using Biopython[15]:
   - Residue solvent accessibility (rsa), secondary structure type (SST categories), backbone angles (phi, psi), distance to ligand, residue and carbon alpha depths.
2. Amino acid indices:
   - Sequence substitution scores from various BLOSUM and PAM matrices[16].
   - Physicochemical indices from AAindex[17] (e.g., hydrophobicity, polarity, molecular volume).
   - Indicators for glycine and proline presence.
   - Pharmacophoric counts per residue: hydrophobic, positive, negative, acceptor, donor, aromatic, sulfur-containing, neutral groups.

**Supplementary Note 2. Hyperparameter tuning and final model configuration.**

Hyperparameters were systematically optimized through Bayesian optimization (using the Weights & Biases Sweep API) to maximize the average Pearson correlation on cross-validation sets. The following hyperparameters were explored:

- Hidden channels: [128, 256, 512]
- Learning rate: [0.0005, 0.001]
- Batch size: [16, 32]
- Epochs: [4, 5, 6, 8, 10]
- Dropout rates (layers 1–6): [0.1, 0.2, 0.3, 0.4, 0.5]
- Activation functions: [tanh, relu, leaky_relu] (layer-dependent)
- Optimizer: Adam
- Weight decay: [0.0005, 0.001]
- Learning rate scheduler: ReduceLROnPlateau with patience (3, 5, 10) and factor (0.1, 0.5, 0.8)
- Number of GCN layers: [1, 2, 3]

The optimal hyperparameters selected based on cross-validation performance were:

- Hidden channels: 128
- Learning rate: 0.0005
- Batch size: 16
- Epochs: 4
- Dropout rates: layer 1 (0.1), layer 2 (0.5), layer 3 (0.1), layer 4 (0.1), layer 5 (0.4), layer 6 (0.5)
- Activation functions: layer 1-3 (tanh), layer 4 (relu), layer 5 (leaky_relu), layer 6 (relu)
- Optimizer: Adam
- Weight decay: 0.001
- Learning rate scheduler factor: 0.8
- Learning rate scheduler patience: 10
- Number of GCN layers: 3

The final model architecture (ProteinLigandGCN) incorporated these hyperparameters during training.
